# Immunogenicity and Structure of a Stabilized HIV-1 clade C Env from a Pediatric Elite Neutralizer in Complex with an Autologous bnAb

**DOI:** 10.1101/2025.04.26.649701

**Authors:** Swarandeep Singh, Sanjeev Kumar, Arnab Chatterjee, Lakshay Malhotra, Muneeb Pervez, Abdul Rahman, Ifrah Andrabi, Bapan Mondal, Sanket Katpara, Shaifali Sharma, Harsh Bhakhri, Anjli Gaur, Tanu Bansal, Vikram Iyer, Abdul Wahid Hussain, Shubbir Ahmed, Rajesh Kumar, Abdul Samath Ethayathulla, Rakesh Lodha, Somnath Dutta, Kalpana Luthra

## Abstract

**Introduction:** The envelope (Env) glycoproteins derived from circulating viruses in elite-neutralizers (ENs), who naturally develop potent broadly neutralizing antibody (bnAb) responses, serve as a potential template for HIV-1 vaccine design. Here, we investigated the immunogenicity and structure of a clade C HIV-1 Env trimer (330) derived from the pediatric elite-neutralizer AIIMS_330.

**Methods:** The 330 Env was engineered as SOSIP.664, NFL.664, and multivalent ferritin-nanoparticle (NP) immunogens. Biophysical and antigenic properties were assessed using HIV-1 trimer-specific bnAbs and non-neutralizing antibodies. Immunogenicity was evaluated in rabbits immunized with Addavax-formulated SOSIP.664, NFL.664, or ferritin NP immunogens (n = 4), with an adjuvant only placebo group (n = 2). Structural characterization was performed by cryo-electron-microscopy (cryo-EM) using the autologous Env 330 SOSIP.664 trimer in complex with the bnAb 44m.

**Results:** The 330 Env trimers exhibited native-like antigenic properties, high thermostability, strong binding to trimer-specific bnAbs, and limited reactivity with non-neutralizing antibodies. Immunization of rabbits elicited strong autologous neutralizing antibody responses together with modest heterologous Tier-1 neutralization. Autologous neutralizing responses were significantly higher for ferritin-NP (P = 0.019) and SOSIP.664 (P = 0.048) than for NFL.664, with no significant difference between ferritin-NP and SOSIP.664 (P = 0.681). A 5.1 Å cryo-EM reconstruction of the autologous Env 330 SOSIP.664–44m complex revealed native-like presentation of the N332-V3 supersite. The structural analysis further suggested the interacting Env contacts E293, R32, K337, and K446, along with glycans N332, N326, N442, and N448 with the bnAb-44m. We identified R327 of the conserved GDIR-motif and the N332 glycan, within the V3-loop, as neutralizing determinants on the 330 Env, with a fold reduction of 116, on testing the mutant 330 viruses with autologous bnAb 44m.

**Discussion:** Pediatric elite-neutralizer AIIMS_330 derived Env trimers represent structurally and antigenically well-characterized clade C immunogens with ability to elicit strong autologous neutralizing antibodies, with modest heterologous Tier-1 neutralization. Together, these findings advance our understanding of clade C Env antigenic variability. Further, the structural and immunological insights support the potential utility of AIIMS_330 derived Env as an antigenic probe and may inform future rational design of Clade C HIV-1 vaccines and structure-guided strategies for eliciting neutralizing antibodies.

## INTRODUCTION

A critical goal in HIV vaccine development is to elicit broadly neutralizing antibodies (bnAbs) in uninfected and high-risk individuals (1). Despite the challenges associated with this goal, two key observations in humans provide a strong rationale for developing vaccines that can elicit bnAbs. First, the identification of elite neutralizers (ENs), a rare group of HIV-1-infected individuals who develop exceptionally potent bnAb responses, has provided valuable insights for rational HIV-1 immunogen design aimed at eliciting similar protective bnAb responses through vaccination (1–3). Second, simian-human immunodeficiency virus (SHIV) challenge studies in nonhuman primates (NHPs) have demonstrated that passive immunization with bnAbs can confer protection against infection, thereby providing strong support for HIV-1 vaccine strategies designed to elicit similar protective bnAb responses in humans following vaccination (4–8). Stabilized HIV-1 envelope (Env) glycoproteins are considered promising candidates for immunogen design to elicit such bnAb responses. The extensively studied clade A pediatric-origin Env BG505 has demonstrated the ability to elicit autologous and, occasionally, Tier-2 heterologous neutralizing responses across multiple animal models and is currently under evaluation in clinical trials (9–17).

It is believed that single Env-based immunization regimens are unlikely to induce such bnAb responses; therefore, multi-stage, multi-component immunization approaches appear to be a promising strategy to guide rare precursor B cells toward bnAb development (18,19). HIV-1 clade C predominates global HIV-1 infection and accounts for approximately 50% of the infections worldwide (20,21). However, a limited number of clade C Envs have been successfully stabilized and characterized as native-like trimers, often with relatively low expression and limited intrinsic stability, thereby posing challenges for the development of clade C Env-based immunogens (22–25). Multivalent display of HIV-1 Env trimers on self-assembling ferritin nanoparticles (NPs) in a repetitive array has proven to be a promising approach, with several studies demonstrating enhanced humoral immune responses and improved immunogenicity in animal models (26,27).

Studies have suggested that HIV-1-infected children who often acquire the infection through vertical transmission develop bnAbs more frequently and earlier than adults (28–33). Such early development of bnAb responses may be attributed to the relative immaturity of the pediatric immune system, in which the founder virus encounters minimal selection pressure compared with adults, thereby giving evolving B cells an advantage in generating neutralizing antibodies against contemporary viruses (28,30). Recent studies suggest that bnAb development is driven by reciprocal virus–host co-evolution, whereby evolving Env variants and the host B-cell repertoire jointly shape the maturation of bnAb responses (34). Characterization of antigenic variability in viral Envs is of particular interest for investigating antigenic determinants associated with bnAb development. Previously, we reported a pair of chronically HIV-1-infected monozygotic twins, AIIMS_330 & AIIMS_329, exhibiting elite plasma neutralizing activity (32). Longitudinal analysis over 5 years revealed sustained evolution of plasma bnAbs in the AIIMS_330 EN, targeting multiple major neutralizing epitopes, including V1V2, V3-glycan, and CD4bs on the HIV-1 Env glycoprotein (30,32,35). Pseudoviral HIV-1 Env clones derived from circulating viruses in the AIIMS_329 and AIIMS_330 pediatric donors exhibited high susceptibility to neutralization by a select panel of HIV-1 bnAbs with diverse epitope specificities, as well as to autologous contemporaneous plasma antibodies (30,32). Env derived from such contemporaneous viruses can serve as valuable templates for HIV-1 immunogen design, while structural characterization of Env trimers in complex with autologous bnAbs from the same donor can provide valuable insights for vaccine development. We recently characterized the antigenicity and immunogenicity of a native-like Env trimer, generated from a virus from AIIMS_329 (25). In earlier studies, we had evaluated the longitudinal antibody response in AIIMS_330 (a pediatric elite neutralizer identical twin of AIIMS_329) (32), and the evolving 44m bnAb lineage (33,36) and defined the interaction of 44m with BG505 Env. The structural, antigenic, and immunogenic properties of the native-like clade C Env, derived from the viruses isolated from AIIMS_330, along with its interaction with the autologous bnAb 44m, remain to be determined, in order to evaluate the potential of the 330 Env to elicit similar responses on immunization.

Here, we designed and characterized an HIV-1 clade C 330 Env trimer derived from a pediatric elite donor, AIIMS_330. Among the longitudinally derived autologous viruses from AIIMS_330, the 330.16.E6 envelope (described as 330 Env in this study) was selected, as it was susceptible to neutralization by multiple bnAbs and contemporaneous autologous plasma antibodies, distinguishing it from several other autologous viral variants that were comparatively resistant, as previously reported (30,32). The 330 Env was engineered as soluble SOSIP.664 and NFL.664 gp140 trimers, and a ferritin NP displaying SOSIP.664 Env trimers in a repetitive array. The designed 330 Envs adopted native-like trimeric conformation and exhibited strong binding to key HIV-1 bnAbs, while showing limited accessibility to the immunodominant epitopes targeted by non-neutralizing antibodies (non-NAbs). The 330 Env constructs elicited strong autologous neutralizing responses, with modest heterologous Tier-1 responses, but did not induce bnAb responses. The multimerization of Env on ferritin NPs resulted in an improved autologous response compared with soluble gp140 trimers. A 5.1 Å cryo-EM reconstruction of the autologous Env 330 SOSIP.664–44m complex revealed a native-like presentation.Collectively, these findings provide mechanistic insights into clade C Env-autologous bnAb interactions and define the structural, antigenic, and immunogenic properties of an Env derived from a pediatric elite neutralizer, while providing a potential antigenic template for future sequential or multicomponent HIV-1 immunogen strategies.

## RESULTS

### 330 Env construct designs and biophysical characterization

We previously reported a pseudoviral Env clone 330.16.E6 derived from an HIV-1 Indian clade C-infected pediatric EN AIIMS_330 that exhibited potent neutralization susceptibility to a select panel of HIV-1 bnAbs targeting V1V2, V3-glycan N332 supersite, CD4bs, gp120-gp41 interface, and MPER epitopes, as well as with autologous contemporaneous plasma antibodies, while showing limited exposure to the panel of non-nAbs tested (30,32). In this study, using 330.16.E6 as a template, we designed and engineered 330 Env constructs in three formats: as soluble cleaved 330 SOSIP.664 gp140 Env trimers, uncleaved 330 NFL.664 gp140 Env trimers, and SOSIP.664 Env trimers displayed on self-assembling 330 ferritin nanoparticles (NPs) to enable multivalent antigen presentation (**Figure 1 A-C, Figure S1**). The 330 Env constructs were stabilized by introducing strategic mutations to increase expression and trimerization, as illustrated in **Figure 1A-D & Table S1**. The codon-optimized Env constructs were expressed in Expi293F cells and initially purified by *Galanthus nivalis* lectin (GNL) affinity chromatography. This initial purification yielded ∼15 mg/L of SOSIP.664 Env and ∼16.5 mg/L of NFL.664 Env, comprising a heterogeneous population of Env oligomers, including monomers, dimers, trimers, and aggregates. To select properly folded Env trimers in the prefusion-closed conformation, Expi293F supernatant was subjected to PGT145-based immunoaffinity chromatography, followed by size-exclusion chromatography (SEC). PGT145 recognizes a quaternary apex epitope preferentially presented by native-like, prefusion-closed Env trimers, thereby enabling enrichment of trimers displaying an intact apex architecture. The resulting PGT145-enriched material was further resolved by SEC to obtain purified Env trimers for subsequent biochemical, biophysical, and antigenic characterization. The SEC profiles of 330 SOSIP.664 and NFL.664 Env trimers showed elution peaks at 10.2 mL volume, whereas ferritin NPs eluted earlier, around 8.7 mL, on a Superdex 200 10/300 column (**Figure 1E**). The purification yields of 330 Env constructs following SEC were ∼1 mg/L for SOSIP.664, ∼1.45 mg/L for NFL.664, and ∼0.05 mg/L for ferritin NPs. Purity analyses confirmed the intact trimeric nature of the gp140 Env trimers, which appeared as crisp bands on Coomassie-stained blue native polyacrylamide gel electrophoresis (BN-PAGE) (**Figure S2A**), whereas the 330 ferritin NPs were too large to efficiently enter the gel and were visible at the top of the lanes. (**Figure S2A, right**). SDS-PAGE analysis showed that the 330 gp140 Env proteins migrated as monomers under reducing conditions, suggesting the high purity of the Env trimer preparations (**Figure S2B**). Negative-stain electron microscopy (ns-EM) reference-free two-dimensional (2D) class averages demonstrated that most of the gp140 proteins adopted a native-like trimeric conformation, with multiple copies of the Env spikes efficiently displayed on the self-assembling ferritin NPs (**Figure 1F and Figure S3**). The 3D reconstruction of soluble gp140 trimers further confirmed a trimeric structure, while the predicted model of ferritin NPs displays multiple copies of SOSIP.664. Env spikes (**Figure 1G**). The thermal stability of 330 SOSIP.664 was evaluated by differential scanning fluorimetry (DSF), which revealed a melting temperature (Tm) of ∼74°C (**Figure 1H**). Overall, biophysical characterization demonstrated that the 330 Envs are well-stabilized and adopt native-like trimeric conformations.

**Figure 1:**
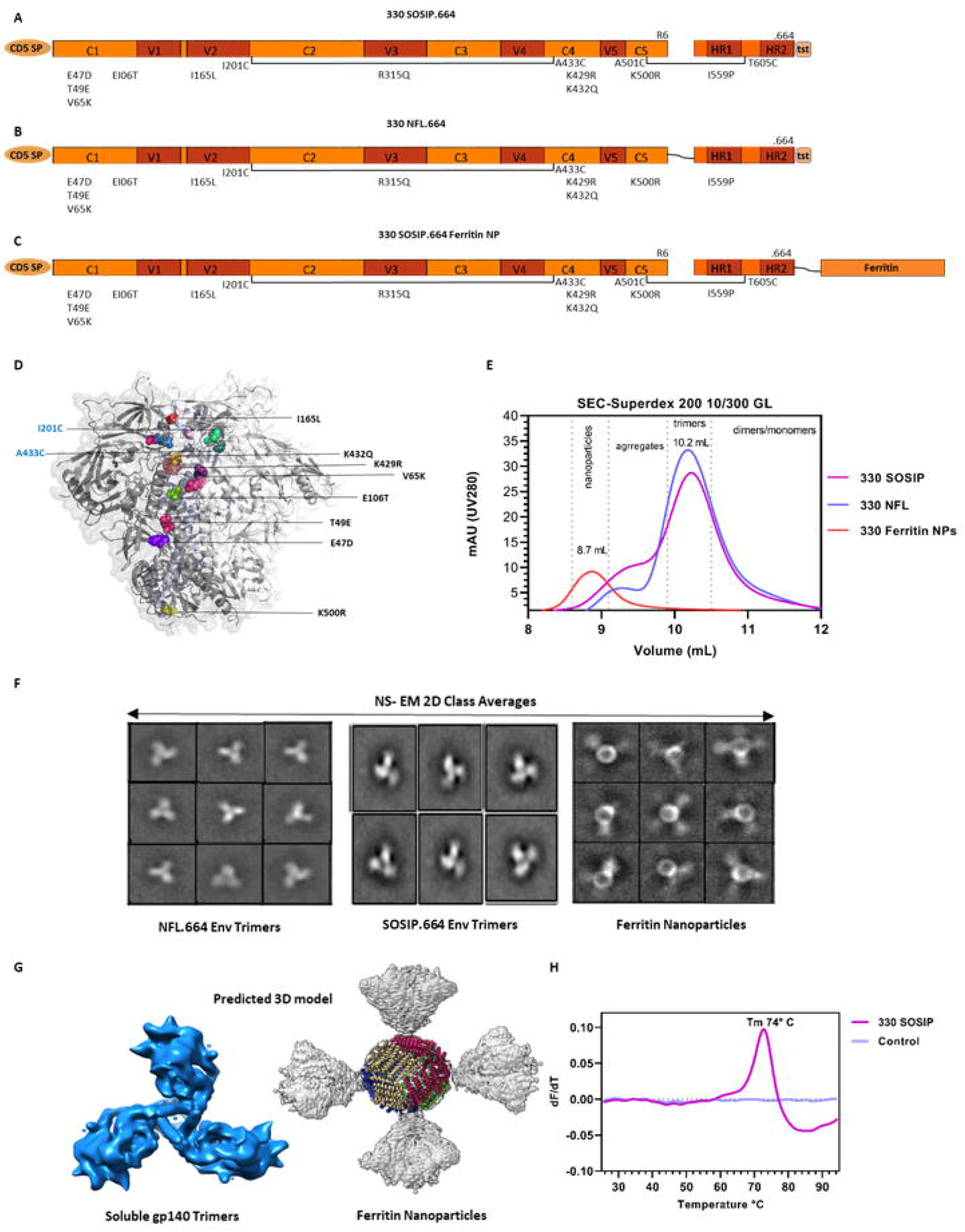
Design and biophysical characterization of 330 gp140 Env trimers and ferritin nanoparticles. **(A-C)** Linear representation of the three 330 Envelope construct design. **(A)** SOSIP.664 design with the SOS-bond (A501C-T605C), IP (I559P) stabilizing mutation along with TD8 stabilizing mutations, an intraprotomer disulfide bond (I201C-A433C), an optimized furin cleavage site (R6), the CD5 Signal Peptide (SP), and Twin Strep-tag (tst). **(B)** NFL.664 design covalently linking gp120 to gp41 with a flexible glycine-serine (GGGGSGGGGS) linker. **(C)** Multivalent design with SOSIP.664 Env displayed on H. pylori ferritin nanoparticle covalently linked with glycine-serine linker at the C-terminal. **(D)** Structural representation of 330 Env highlighting the TD8 stabilizing mutations and I201C-A433C disulfide bond. The structure was illustrated using PyMOL and was based on the BG505 SOSIP.664.664 crystal structure (PDB ID: 4ZMJ). **(E)** SEC profiles of 330 Env trimers (SOSIP.664 and NFL.664) eluted around 10.2 mL, and Ferritin NP showed a major peak at 8.7 mL, using a Superdex 200 Increase 10/300 GL column. **(F)** Negative stain-electron microscopy (ns-EM) 2-D class averages of NFL.664 and SOSIP.664 gp140 Env trimers and ferritin NP. **(G)** Predicted 3D model of Env trimers and ferritin NP displaying SOSIP.664 Env trimers using ns-EM 2D class averages using ChimeraX. **(H)** Thermal stability profile of 330 SOSIP.664 Env trimer determined by DSF.

### 330 gp140 Env trimers and ferritin nanoparticles exhibit strong binding to HIV-1 bnAbs

Next, we evaluated the antigenic profiles of the three 330 Envs (SOSIP.664, NFL.664, and ferritin NPs) by ELISA assays using a select panel of HIV-1 bnAbs including those targeting the V1V2 apex (PGT145, PGDM1400, CAP256.25), the V3-glycan (2G12, PGT121, 10-1074, AIIMS P01), the CD4 binding site (VRC01, N6), and the gp120-gp41 interface (PGT151). We also evaluated the binding of Envs with a panel of non-nAbs, including F105, which targets CD4bs on open Env conformations, A32, which recognizes the gp120 inner domain for open states, and the CD4-induced antibodies 48d and 17b, assessed in the presence of soluble CD4 (sCD4) (**Figure 2**). All the three 330 Env constructs were recognized by the tested HIV-1 bnAbs, with the exception of PGT151, which did not bind the uncleaved NFL.664 trimer, consistent with the absence of the native gp120–gp41 cleavage-dependent interface required for PGT151 recognition. In contrast, the constructs showed minimal binding reactivity with the CD4-induced non-neutralizing antibody 17b in the presence of sCD4, supporting their predominantly closed, native-like prefusion conformation. Overall, reactivity to the non-neutralizing antibodies was lower than that observed for broadly neutralizing antibodies, although some differences in epitope exposure were evident among the platforms. For F105, NFL, and ferritin NP, AUCs were modestly higher than SOSIP, at 2.43, 3.00, and 1.78, corresponding to 1.37- and 1.69-fold increases, respectively. Similarly, A32 binding increased 1.44-fold for NFL (AUC = 2.39) and 1.27-fold for ferritin NP (AUC = 2.11) compared with SOSIP (AUC = 1.66). For 48d, reactivity was modestly higher with NFL (1.23-fold; AUC = 2.31) and ferritin NP (1.45-fold; AUC = 2.71) than with SOSIP (AUC = 1.88). In contrast, 17b binding in the presence of sCD4 showed a marked increase for NFL, with a 3.22-fold higher AUC (3.41) than SOSIP (1.06), whereas ferritin NP showed only a modest 1.35-fold increase (AUC = 1.43). Overall, the 330 gp140 SOSIP, NFL, and ferritin NP platforms showed high binding reactivity with the bnAbs tested, with negligible binding to non-neutralizing antibodies.

**Figure 2:**
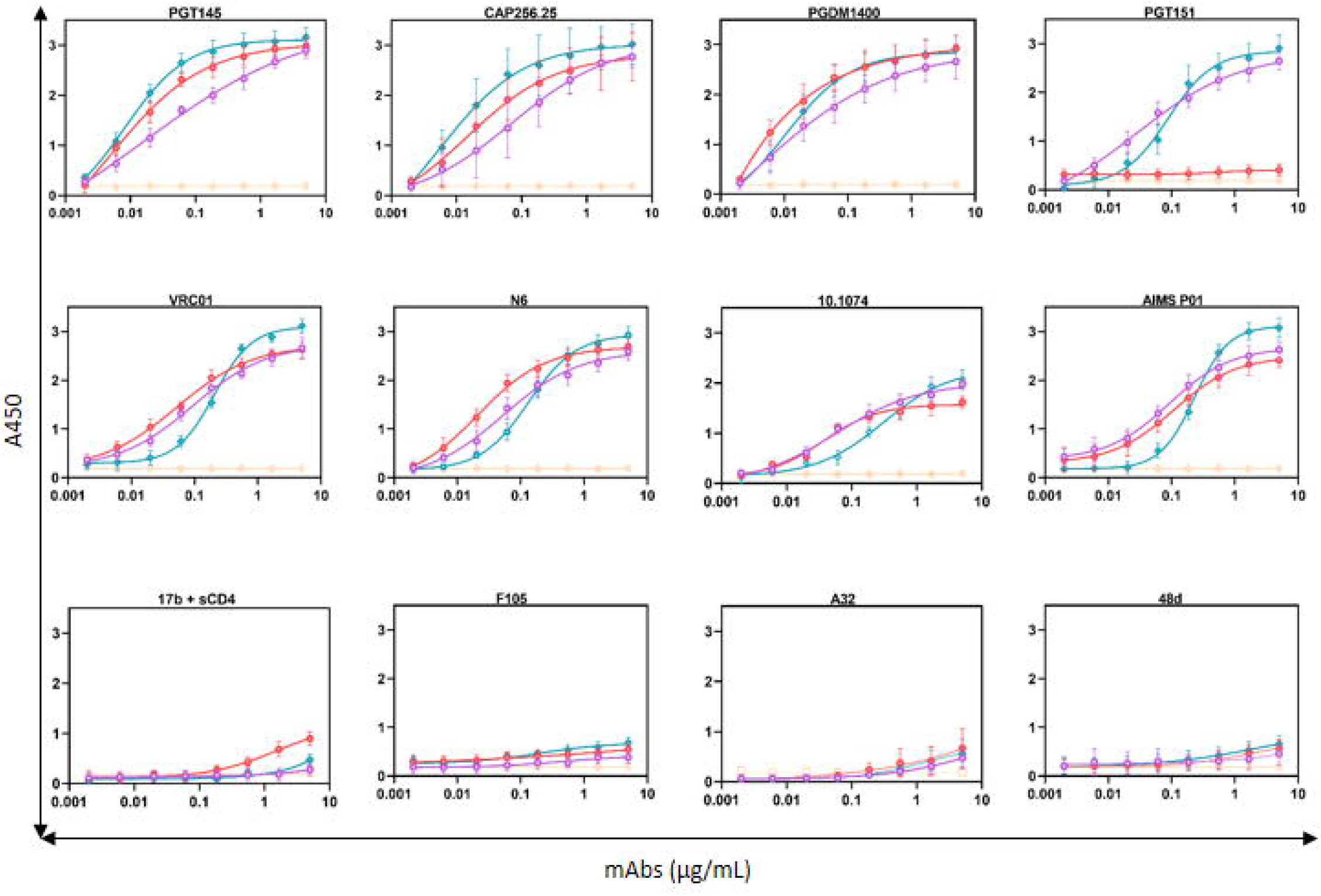
Antigenicity of 330 gp140 Env trimers and ferritin nanoparticles to HIV-1 broadly neutralizing and non-neutralizing antibodies. ELISA binding curves showing the antigenicity of 330 Env trimers and ferritin nanoparticles with conformation-specific bnAbs, including V1V2-directed (PGT145, CAP256.25, PGDM1400), gp120-gp41 interface (PGT151), CD4bs (VRC01, N6), V3 Supersite directed (10-1074, AIIMS P01), along with non-nAbs, including CD4-induced co-receptor binding site (CD4i) non-nAbs (17b+sCD4, A32, 48d) and CD4bs non-nAb (F105). Data are presented as mean values ± SD from two independent experiments, each performed in duplicate.

Subsequently, we evaluated the binding kinetics of all three 330 Env constructs using octet biolayer interferometry (BLI) analysis. We used selected HIV-1 trimer conformation-specific bnAbs PGT145, PGDM1400, and PGT151, along with non-nAbs F105 and 17b, in the presence of sCD4 (Figure 3). Sensograms were fitted using a 1:1 binding model, and the resulting kinetic parameters are reported as KD for SOSIP.664 & NFL.664 Env trimers and apparent dissociation constants (*K*_D_^app^) for Ferritin NPs, as multivalent interactions may introduce avidity and rebinding effects that can influence the observed binding kinetics. The 330 Envs SOSIP.664 and NFL.664 exhibited comparable binding to the V2-apex-directed bnAb PGT145, with KD values of 1.07 nM and 1.39 nM and a *K*_D_^app^ of 1.10 nM for ferritin NP, respectively (**Figure S4**). A similar binding affinity pattern was observed for PGDM1400, with a KD of 1.34 nM for 330-SOSIP.664, KD of 3.32 nM for the 330-NFL.664 Env and *K*_D_^app^ of 1.39 nM for the ferritin NPs. An HIV-1 cleaved trimer and gp120-gp41 interface-directed bnAb PGT151 showed strong affinity to both 330-SOSIP.664 and ferritin NPs, with KD of 5.0 pM and *K*_D_^app^ of 3.0 pM, while no binding was observed to NFL.664, which is an uncleaved trimer design. Consistent with the ELISA data, all three Envs exhibited very low binding affinity to non-nAbs (**Figure S4 & Table S2**). Collectively, the antigenic characterization of all three 330 Envs confirmed their high binding and affinity to key bnAbs and limited recognition by immunodominant non-nAbs.

**Figure 3:**
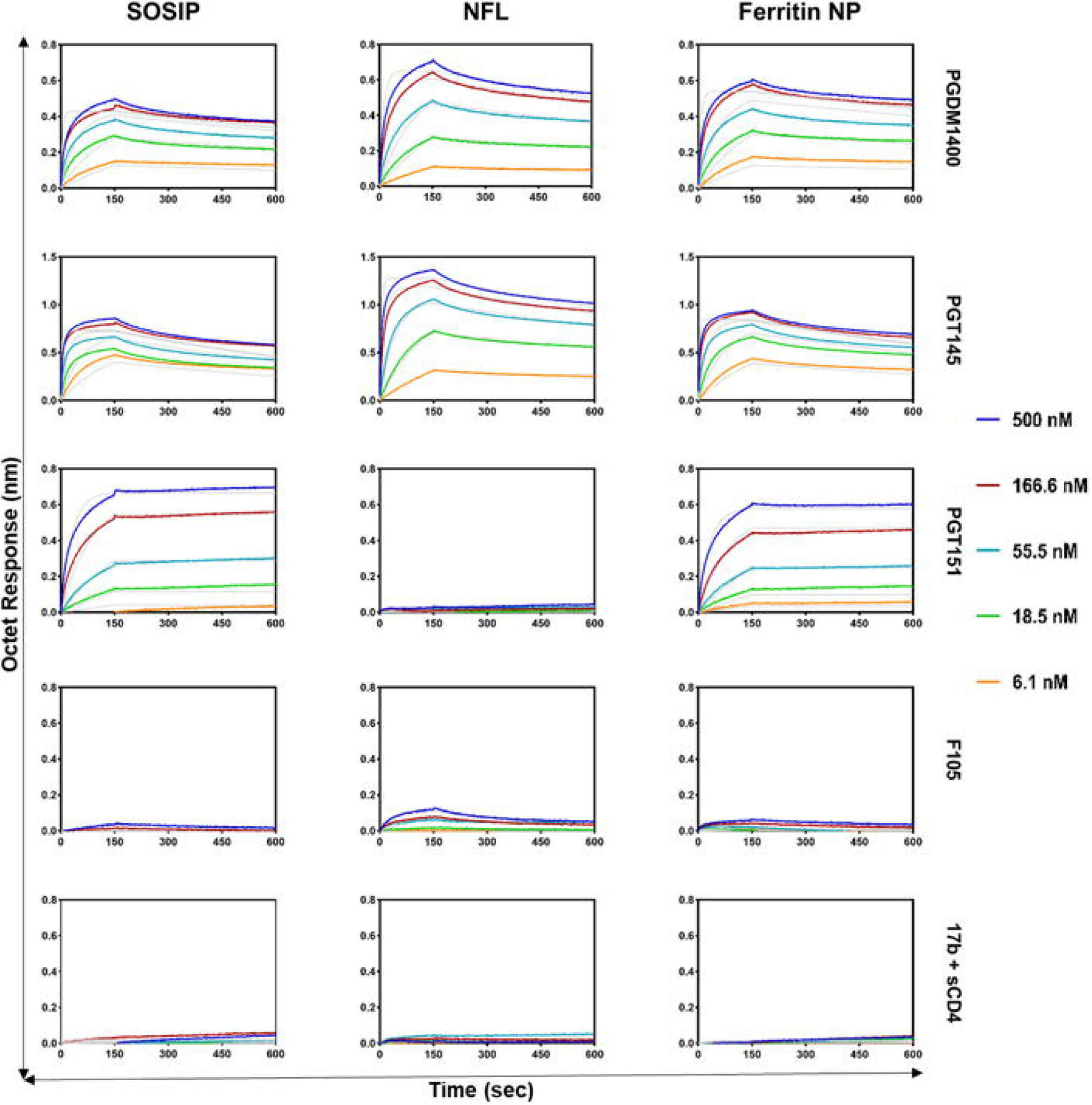
BLI-Octet binding kinetics of 330 Env with HIV-1 broadly neutralizing and non-neutralizing antibodies. BLI Octet sensorgrams showing response shift (nm) on the y-axis against over time (sec) at the x-axis of 330 gp140 Env trimers and NP with varied concentrations (500 nM, 166.6 nM, 55.5 nM, 18.5 nM, 6.1 nM) against the quaternary specific bnAbs PGT145, PGDM1400, cleavage dependent bnAb PGT151, and the non-nAbs F105 & 17b+sCD4.

### 330 SOSIP.664 Env trimer and ferritin nanoparticle elicit potent autologous neutralizing antibody responses compared to NFL.664 trimer in rabbits

Given their favorable antigenic profiles to key HIV-1 bnAbs and native-like conformations, we next evaluated and compared the immunogenicity of soluble 330 Env trimers (SOSIP.664 and NFL.664) and multivalent ferritin NP in New Zealand white rabbits. Four groups of female rabbits were immunized with 20 µg of 330 gp140 Env trimers or an equimolar amount (25 µg) of ferritin NP, formulated with AddaVax adjuvant. Rabbits were primed at week 0, and the first boost or second dose was administered at week 4, and the second boost or third dose was administered at week 20. (**Figure 4A**). The AddaVax is a squalene-based oil-in-water nanoemulsion that enhances vaccine immunogenicity by promoting both cellular (Th1-associated) and humoral (Th2-associated) immune responses and has previously been shown to preserve the structural integrity and antigenicity of the HIV-1 Env trimer (37,38). Rabbits were bled at weeks 0, 4, 6, 8, 12, 16, 20, and 22, and sera samples were subsequently evaluated for binding reactivity with the Env trimer using a capture ELISA and for neutralizing activity against a select panel of Env-pseudotyped viruses. First, we evaluated the binding titers of the collected immune sera at different timepoints against the respective immunogens and determined the median titers for each group. Serum binding titers increased markedly between weeks 4 and 8 and again between weeks 20 and 22, corresponding to the second and third boosters (**Figure 4B**). The polyclonal immune sera exhibited sustained binding reactivity against the autologous Env trimer as well as cross-clade heterologous Envs, although responses to the heterologous Envs were comparatively lower (**Figure 4C**). Using the autologous 330 Env, we observed an evolving polyclonal immune sera binding response following the first boost, with higher binding reactivity with ferritin NP (1.539 ± 0.151) and SOSIP (1.453 ± 0.147) than NFL (1.209 ± 0.059) (**Table S10**). Tukey’s HSD analysis confirmed significant differences between ferritin NP and NFL (P = 0.019) and SOSIP and NFL (P = 0.048), but not between ferritin NP and SOSIP (P = 0.681). At Week 20, binding reactivity to ferritin NP remained significantly higher than SOSIP (P = 0.035) and NFL (P = 0.041), with relatively higher binding to SOSIP than NFL (P = 0.002). Following recall at Week 22, the immune sera showed significantly higher binding with ferritin NP than to both SOSIP (P = 0.002) and NFL (P = 0.001). Binding of the immune sera with SF162 recombinant protein showed no significant differences at weeks 6, 20, or 22 (P > 0.05). On testing the ferritin NP immune sera, binding reactivity with the 6240 gp140, showed higher week 6 binding (1.524 ± 0.085) than that of SOSIP (1.334 ± 0.050; P = 0.015), whereas the remaining pairwise comparisons at Weeks 20, and 22 were not significant (P > 0.05). Binding reactivity of ferritin NP immune sera, with the 1086 gp140, showed that week 6 binding was higher (1.067 ± 0.104) than that of the NFL (0.646 ± 0.027) and SOSIP (0.091 ± 0.174) immune sera, with significant differences for ferritin NP versus NFL (P = 0.002) and SOSIP versus NFL (P = 0.026). The binding kinetics for the ferritin NP immune sera with 329 gp140 (1.458 ± 0.115) was lower than of SOSIP (1.672 ± 0.107; P = 0.031) and, NFL (1.626 ± 0.062). Overall, the strongest and most consistent immune sera reactivity was observed against the autologous 330 gp140 than the heterologous Env proteins.

**Figure 4:**
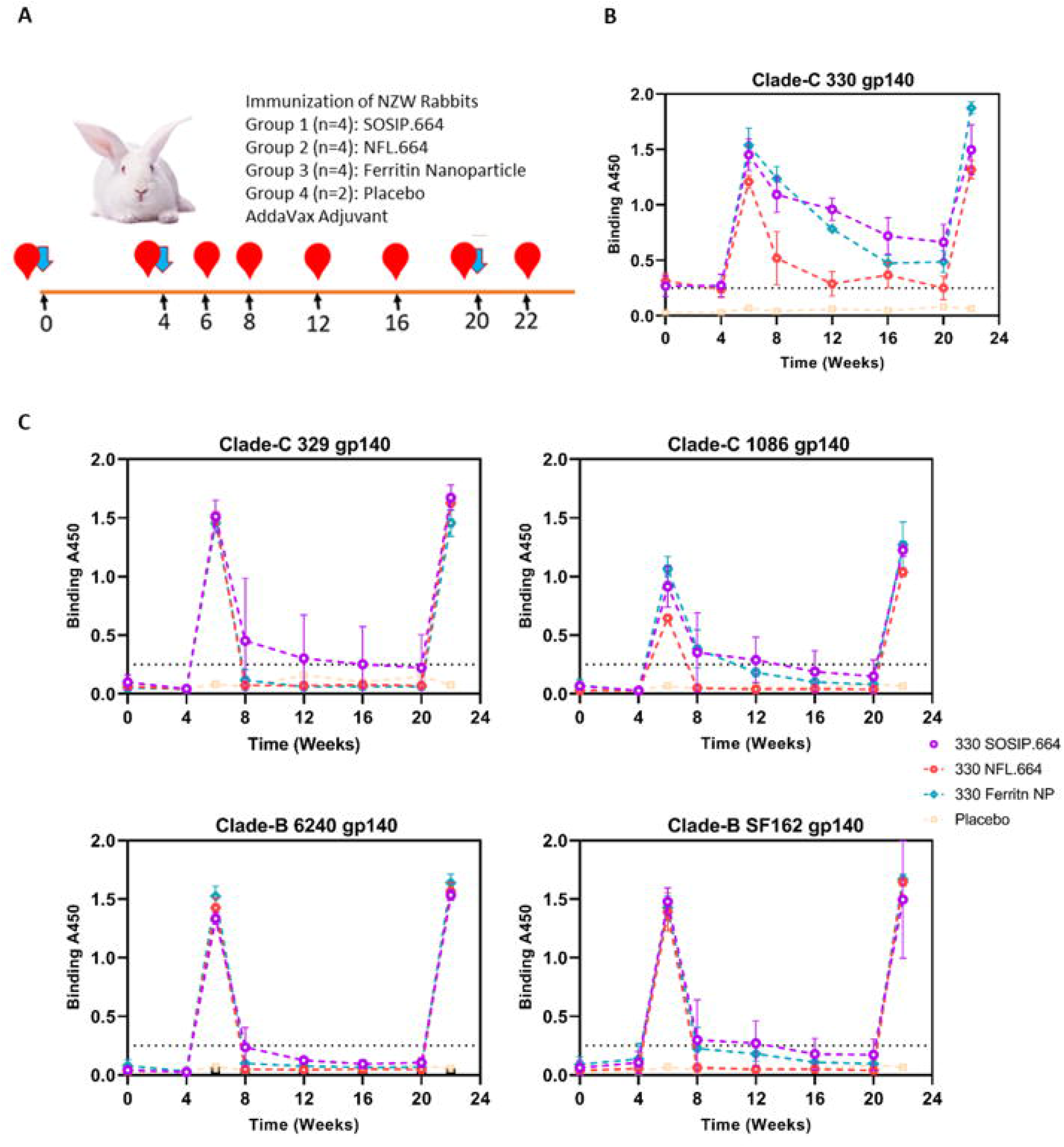
Immunogenicity of 330 SOSIP.664, NFL.664 Env trimers & Ferritin NPs in NZW rabbits. **(A)** Rabbit immunization Schedule. Four groups of rabbits were primed at week 0 and boosted at weeks 4 & 20 (blue arrows). Group 1 immunized with 20 µg SOSIP.664 Env gp140 trimers (n=4), Group 2 immunized with 20 µg NFL Env gp140 trimers (n=4), Group 3 immunized with 25 µg Ferritin Nanoparticles (n=4), and Group 4 served as Placebo (n=2), received AddaVax adjuvant (1:1 v/v) formulated with sterile 1X PBS. Serum samples were collected at weeks 0, 4, 6, 8, 12, 16, 20 & 22 (red arrows). **(B)** Longitudinal binding (OD at A450) against clade-C autologous Env 330 and against the heterologous Envs, including Clade-C 329 and 1086, Clade-B Envs SF162 and 6240. Longitudinal serum-binding responses were analyzed using linear mixed-effects models (LMMs) to account for repeated measurements within animals and to assess the effects of immunogen group, time, and group × time interaction. Statistical significance was defined as **P** < 0.05, and exact **P** values are reported where applicable. Data are presented as mean ± standard deviation (SD).

The peak serum binding response to 330 Envs was observed at week 22, and immune sera were subsequently tested to evaluate neutralizing activity. Neutralization breadth of the immune sera was evaluated against autologous pseudovirus (330.16.E6), closely related pseudovirus (329.14.B1), and Tier 1 heterologous clade C (MW965.26 and 25925), and clade B (SF162 and Bal01) pseudoviruses. All animals exhibited neutralizing antibody responses following immunization, except for one in the NFL.664 group (NZW5273) (**Table S5**). No neutralizing activity was detected against murine leukemia virus (MLV) used as a negative control. Multivalent 330 ferritin NPs and SOSIP.664 elicited a strong nAb response against autologous (330.16.E6) and closely related Env (329.14.B1) pseudoviruses, as compared to NFL.664 trimer (**Figure 5A**). The ferritin NP group elicited the strongest autologous neutralizing antibody response, with a median ID_50_ titer of 1160, compared with the soluble gp140 SOSIP.664 and NFL.664 trimers, which had median ID_50_ titers of 830 and 375, respectively. The nanoparticle group elicited significantly higher nAb titers than both the SOSIP.664 (P = 0.01) and NFL.664 (P = 0.01) groups, with fold increases of 1.4 and 3.1, respectively. These findings were consistent with the immune sera binding profiles, in which the ferritin NP and SOSIP.664 groups maintained sustained binding reactivity, whereas the NFL.664 group exhibited a waning of antibody responses following the early post-immunization responses, but rose again at the week 20 boost. The ferritin NP and SOSIP.664 groups elicited moderate heterologous nAb responses against the clade C Tier-1 viruses MW965.26 and 25925, as well as the clade B Tier 1 virus SF162, with comparable responses between the two groups and higher titers than those observed in the NFL.664 group, although these differences did not reach statistical significance (**Figure 5B**). In contrast, none of the 330 immunogens induced detectable nAb responses against the clade B Tier 1 virus BaL.01, the clade A Tier 2 virus BG505, indicating limited cross-reactive Tier-1 and Tier-2 heterologous neutralization (**Table S5**). Collectively, the 330 immunogens elicited predominantly autologous neutralizing responses, with modest cross-clade neutralization breadth. Next, to map the Env regions targeted by 330 Env elicited antibodies in NZW rabbits, serum competition analysis was performed using BLI-based tandem epitope binning. Polyclonal immune sera were assessed for competition against a panel of HIV-1 bnAbs, including V1V2 apex-directed (PGT145 and PGDM1400), CD4bs-directed (VRC01), V3-glycan supersite-directed (44m, 10-1074, and PGT121), and the gp120-gp41 interface-directed (PGT151), along with non-nAbs (F105, 447-52D, 412d, and 17b). Rabbit serum competition mapping suggested that immunization elicited polyclonal antibodies against overlapping antigenic regions of the native-like 330 Env trimer, including the V1V2 apex, CD4bs, and V3-glycan supersite (**Figure 5C, Table S6**). However, these competition profiles do not establish that the immunizations elicited antibodies reproduce the V1V2, CD4bs, or V3-glycan-directed bnAb specificities.

**Figure 5:**
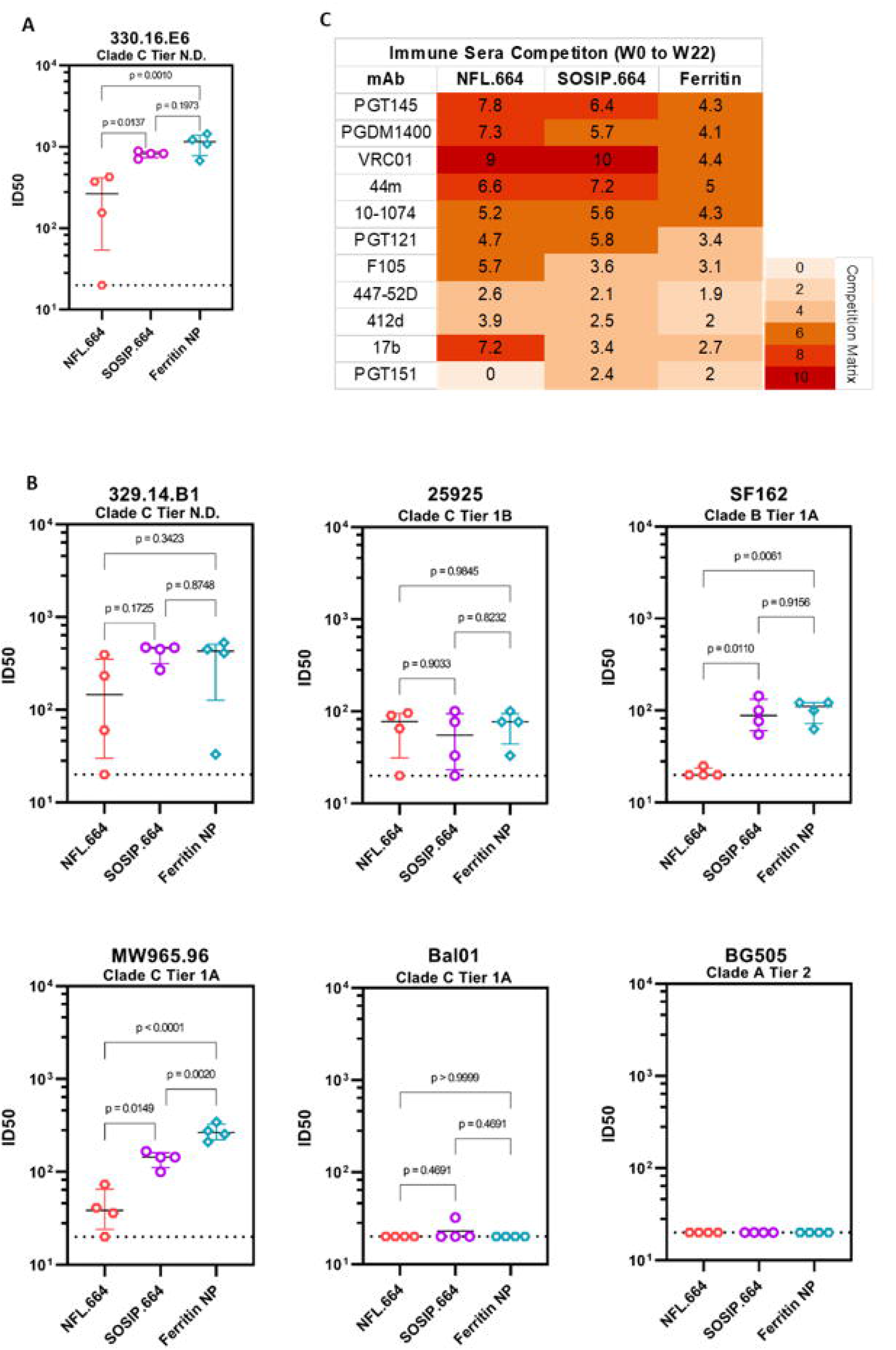
Neutralizing potential of 330 SOSIP.664, NFL.664 Env trimers & Ferritin NPs in NZW rabbits. **(A)** Midpoint neutralization titers (ID_50_) for week 22 sera of the immunized rabbits against autologous pseudovirus 330.16.E6 and fold difference between SOSIP.664, NFL.664 & Ferritin NPs **(B)** ID_50_ titers against closely related pseudovirus 329.14.B1 and heterologous Tier 1 clade-C pseudovirus MW965.26 and 25925, clade-B pseudovirus SF162 and Bal.01, and against Tier 2 Clade-A BG505 pseudovirus. Median ID_50_ titers in the NFL.664, SOSIP.664 & Ferritin NP groups were 375, 830 & 1160, respectively. **(C)** BLI-Octet tandem competition matrix illustrating competitive binding between immune sera and reference HIV-1 bnAbs, including V1V2 apex-directed (PGT145 and PGDM1400), CD4bs-directed (VRC01), V3-glycan supersite-directed (44m, 10-1074, and PGT121), and the gp120-gp41 interface-directed (PGT151), along with non-nAbs (F105, 447-52D, 412d, and 17b). A 35% competition threshold was used to define competing interactions. Competition profiles were analyzed by calculating the percent inhibition of monoclonal antibody binding in the presence of immune sera. ELISA and neutralization assays were performed in duplicate, and data are representative of three independent experiments. The assay cut-off at ID50 titer of 20 is indicated by a dotted line. Individual data points are shown as scatter dot plots, with the median and interquartile range (IQR) as indicated by colored lines. Differences among the SOSIP.664, NFL.664, and Ferritin NP groups were evaluated using unpaired two-tailed **t**-tests or one-way ANOVA followed by Tukey’s multiple-comparisons test, as appropriate. A P value <0.05 was considered statistically significant, and exact P values are shown in the figure.

### Structural and glycan prediction analysis of HIV-1 clade C 330 Env

Next, to investigate the preservation of native Env conformation, we determined the structure of the unliganded 330 SOSIP.664 trimer by single-particle cryo-electron microscopy (cryo-EM) (**Figure S6 & S7)**. A total of 89209 particles were analyzed to determine 2D class averages and to reconstruct the trimeric 3D model at 7.43 Å resolution, suggesting that the 330 SOSIP.664 Env adopted prefusion-closed conformations (**Figure S6 & Table S7**). Further to assess structural similarity, the HIV-1 clade C 330 SOSIP.664 trimer was superimposed onto a clade C Env 16055 NFL.664 (39) (PDB: 5UM8) and a clade-A Env BG505 SOSIP.664 (40) (PDB: 4TVP). We observed that the 330 SOSIP.664 Env trimer exhibited structural similarity to BG505 Env 16055 Env (**Figure 6A-D**). In a glycan prediction analysis using the Los Alamos HIV-1 sequence database (LANGT), we observed that 330 Env lacked conserved PNGS at positions 295 and 339 in the V3/C4 region and at position 130 in the V1 apex region of the envelope (**Figure 6E**). Using the Los Alamos Glycan Shield Mapping tool (LAGSM), we predicted the total glycan hole area of the 330 Env and observed a moderate protein-accessible surface area of 967 Å², which was lower than those estimated for BG505 (2591.12 Å²), B41 (1649 Å²), and C97ZA (1027.09 Å²), but higher than that estimated for Du422 (239.28 Å²) (**Figure 6F**). The potential N-glycosylation sites (PNGS) conserved in the 330 Env were determined by LANGT using the N-Glycosite tool. Conserved PNGS prediction analysis revealed that 330 Env comprises 28 PNGS sites, comparable to those reported for the BG505 (41) and 16055 (42) Envs, whereas B41, C97ZA, and Du422 were predicted to contain 29-30 PNGS sites (**Figure 6F, Table S3**). Collectively, structural homology modeling and glycan shield analyses demonstrated that the stabilized clade C 330 Env shares several structural and glycan features with the well-characterized BG505 Env, supporting its ability to adopt a native-like and well-ordered trimeric conformation.

**Figure 6:**
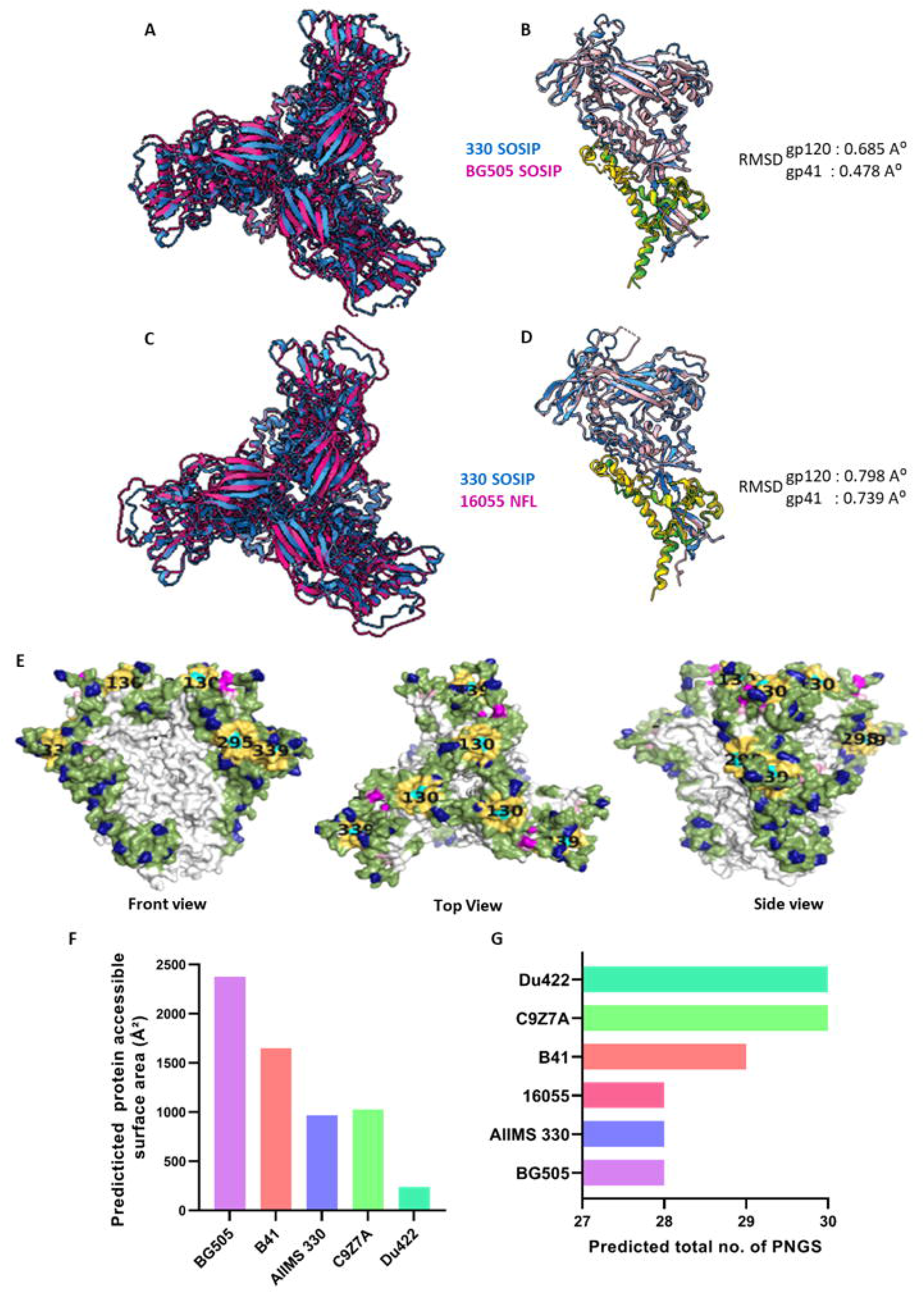
Structural and glycan prediction analysis of 330 SOSIP.664 with BG505 & 16055 trimers. **(A–B).** Structural superposition of the unliganded 330 SOSIP.664 Env trimer (PDB: 9XC1) with the BG505 SOSIP.664 trimer (PDB: 4TVP), illustrated as a trimer and protomer overlay. Root-Mean-Square Deviation (RMSD) values between superimposed protomers were 0.685 Å for gp120 and 0.478 Å for gp41. **(C–D)** Structural comparison of the unliganded 330 SOSIP.664 with the 16055 NFL.664 Env trimer (PDB: 5UM8), with RMSD values between aligned protomers of 0.798 Å for gp120 and 0.739 Å for gp41. Segmented EM densities are colored as follows: 330 SOSIP.664-dodger blue; BG505 SOSIP.664 and 16055-deep pink. Within each protomer, gp120 regions are shown in blue or light pink, whereas gp41 regions are shown in yellow or green. **(E)** Surface representation of predicted glycan holes in the 330 Env trimer at positions 130, 295, and 339 (HXB2 numbering), shown in top, front, and side views. The glycan prediction analysis was performed using the Los Alamos HIV-1 sequence database (LANGT) **(F).** Comparison of predicted total protein-accessible surface area across cross-clade HIV-1 Env trimers was predicted using the Los Alamos Glycan Shield Mapping tool (LAGSM) **(G).** Prediction and Comparison of potential N-linked glycosylation sites (PNGS) between 330 and BG505 Env sequences. “N” indicates the presence of a PNGS motif (N-X-T/S, where X is any residue except proline). Glycan annotations were generated using the LANGT N-Glycosite tool.

### Cryo-EM structure of the autologous 330 Env-44m bnAb complex reveals GDIR and N332 dependence

To further confirm the structural integrity and antigenic recognition of 330 SOSIP.664 Env by its autologous HIV-1 bnAb, we next performed its structural analysis in complex with a heavy-chain-matured HIV-1 bnAb 44m. The 44m bnAb is a lineage member of the AIIMS 330 pediatric elite neutralizer-derived bnAb AIIMS P01 (30,36), which potently neutralizes its autologous pseudovirus, 330.16.E6, from which the 330 Env trimer is derived (33). We determined the cryo-EM structure of the 330 SOSIP.664 Env trimer in complex with the donor-matched 44m Fab at 5.1 Å resolution (**Figure S8, Table S8**). The 330 Env adopted a native-like HIV-1 Env trimer conformation, with three additional densities corresponding to bound 44m Fab molecules clearly visible on the trimer surface. (**Figure 7A-B, Figure S8**). The model revealed that one 44m Fab engaged each gp120 protomer within the trimer, resulting in a 3:1 Fab-to-trimer ratio, suggesting efficient trimer engagement (**Figure 7C-D**). The interacting surface areas contributed by the 44m antibody and the 330 Env were approximately 690 Å² and 613 Å², respectively. Structural analysis further revealed that the CDRH1 loop of 44m was positioned proximal to the V1V2 region of the 330 Env, whereas the CDRH2 and CDRH3 loops primarily engaged the V3-glycan supersite (**Figure 7E**). The autologous 330 Env-44m complex showed dependence on R327 of the conserved GDIR motif and the N332 supersite, as assessed by mutant viral neutralization assays. Structural analysis further suggested Env contacts involving residues E293, K337, and K446, along with glycans N326, N442, and N448 with the bnAb 44m. (**Figure 7F & H**).. Distinct contact residues identified in the autologous 330 Env-44m complex included S30 in the CDRH1 loop interacting with R327 in the V3 region; Y52 and Y53 in the CDRH2 loop contacting R327 and H330; and R109 in the CDRH3 loop engaging E293 in the C2 region, K337 in the C3 region, and K446 in the C4 region of gp120 (**Figure 7F**). In addition, the R78 residue within the FR3 region formed interactions with D325 in the GDIR motif, indicating that both the CDRH2 and FR3 regions contribute to stabilization of the Env-antibody interface (**Figure 7E-F**). The observed interactions are predicted to contribute to the stabilization of the Env–44m bnAb interface through hydrogen bonding, electrostatic interactions, and van der Waals forces. Electrostatic surface potential analysis demonstrated that residues E293, D325, R327, H330, N332, K337, and K446 were oriented toward charged and hydrophilic regions within the 44m paratope (**Figure 7G**). Glycan-mediated interactions at the 330 Env-44m interface involved N301 and N332 glycans within the V3 glycan supersite, together with N397 in the V4 region and N442 and N448 within the C4 region (**Figure 7H**). Collectively, these findings demonstrate that the autologous 330 Env engages 44m through an expanded peptide-glycan interaction network relative to BG505, highlighting the structural adaptability of V3-glycan supersite recognition during autologous Env-antibody co-evolution.

**Figure 7:**
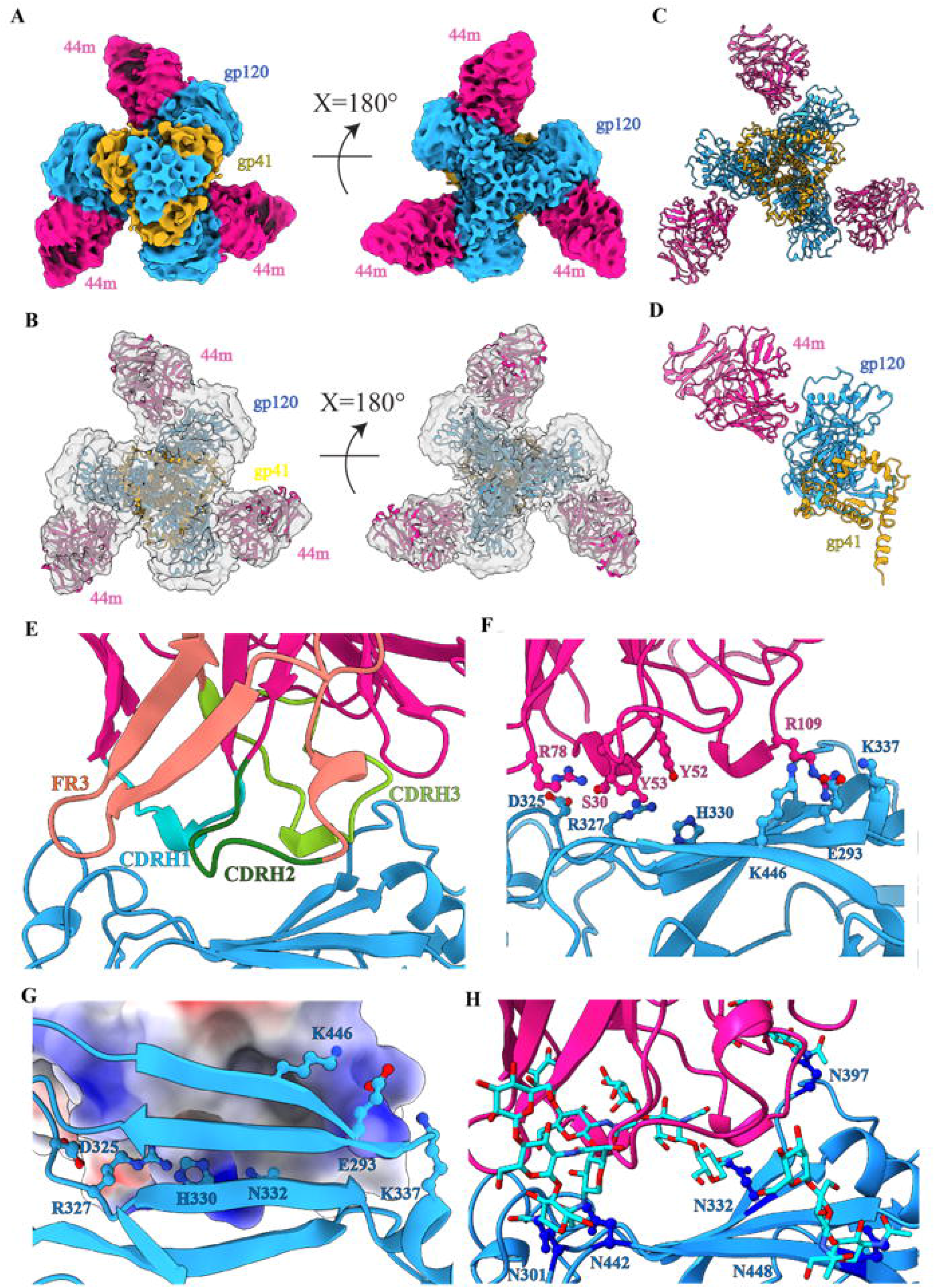
Cryo-EM structure of 330 SOSIP.664 Env trimer in complex with autologous bnAb 44m. **(A)** Top and side views of cryo-EM reconstructed map of 330 SOSIP.664 Env trimer in complex with 44m bnAb. Color coding corresponding to segmented EM densities is: 44m bnAb-deep pink, gp120-dodger blue, gp41-golden rod; **(B)** The corresponding atomic model fitted in the EM map shown in grey. **(C)** Structural model illustrating binding three Fabs of 44m to the trimeric 330 Env complex, with one Fab engaging each gp120 protomer **(D)** Enlarged view of a single 330 protomer in complex with one 44m Fab **(E)** Localization of 44m CDRH and framework regions (FR) involved in Env recognition, colored as: CDRH1-cyan, CDRH2-deep green, CDRH3-light green, FR3-light brown **(F)** Key interacting residues 44m bnAb - S30, Y52, Y53, R78 and R109 and corresponding interacting residues of gp120 loops - E293, D325, R327, H330, N332, K337 and K446 highlighted in heteroatom color. The distances between S30 and R327; Y52 and H330; Y53 and R327; R78 and D325 are 3.4 Å, 5.6 Å, 3.8 Å, 6.3 Å, respectively. The separations between R109 and E293, K337, and K446 are 3.6 Å, 3 Å, and 6.5 Å, respectively. **(G)** Electrostatic surface potential map illustrating charge complementarity between gp120 interface residues and the 44m paratope. **(H)** N-linked glycans N301, N332, N397, N442, and N448 are located at the 330-44m interface.

## DISCUSSION

HIV-1 clade C accounts for more than half of global HIV-1 infections, with particularly high prevalence in South Africa and India (20,21). Comparative analyses of clade C viruses from these regions reveal the significant differences in their sequences, highlighting distinct evolutionary trajectories (43). These regional variations in intra-clade C Env sequences likely contribute to the differential sensitivity of circulating strains to HIV-1 nAbs. HIV-1 clade C is characterized by unique features such as enhanced transmission efficiency, extensive genetic diversity within the V1-V3 region of the Env gene, and a distinct profile of drug resistance mutations compared to other clades (43–46). Understanding these differences is crucial for developing prevention and treatment strategies in regions where clade C is prevalent (20,21). One of the most extensively studied and characterized HIV-1 Env immunogens, BG505, is a clade A transmitted/founder Env derived from a 6-week-old infant with elite neutralizing activity (9). Such elite individuals from the pediatric group could provide valuable Env immunogen templates, as HIV-1 infection and disease progression in children differ from that in adults (28,31,32). Notably, bnAbs tend to arise early in the course of infection in HIV-1-infected children (30,47). From the pediatric cohort, we previously reported a pair of monozygotic twins (AIIMS_329 and AIIMS_330) with elite neutralizing activity against contemporaneous viral Env (30,32).

Here, we describe the development and characterization of native-like clade C 330 Env trimers derived from pediatric EN AIIMS_330, generated in both SOSIP.664 and NFL.664 formats, as well as the multivalent presentation of the SOSIP.664 Env on self-assembling ferritin NPs. The Negative stain-Electron microscopy (ns-EM) analysis showed 330 Env constructs adopted key features of native-like trimeric conformation, and Env trimers were efficiently displayed on ferritin NP, consistent with previous observation for stabilized BG505 Env-particles (26). The purification yields of PGT145-purified 330 Env trimers following SEC were ∼1 mg/L for the SOSIP.664 construct and ∼1.45 mg/L for the NFL.664 construct. The yields were comparable to or higher than those for previously reported Indian and South African clade C Env trimer, including CZA97: ∼0.4-0.6 mg/L (48), 329 SOSIP.664: ∼0.6 mg/L (49), although lower than the yield reported for the 16055 NFL.664: ∼2 mg/L (50). Biophysical characterization showed 330 Env exhibits a high degree of thermal stability with a melting temperature (Tm) of ∼74°C, which is higher than that of other reported clade C Envs, including DU422 (63.4°C)(51), ZM197 (62.7°C)(51), and 329 SOSIP.664 (68.1°C) (49), IPGE-THIVC (62°C)(24), and remaining lower than that reported for 16055 (77°C) (50). The antigenic profiling of the 330 Envs demonstrated nanomolar binding to conformation-dependent bnAbs, indicating proper presentation and accessibility to bnAb epitopes. In contrast, minimal binding reactivity to the non-nAbs tested suggested effective shielding of immunodominant non-neutralizing regions. The incorporation of the previously described 201C-433C intraprotomer disulfide bond (52) likely restricted the CD4-induced Env conformational rearrangements, thereby stabilizing the Env proteins in a closed prefusion conformation.

Previous studies have established rabbits as a widely used preclinical model for evaluating the immunogenicity of HIV-1 Env immunogens before their assessment in NHP models and clinical studies (53,54). Moreover, comparative analyses have shown that vaccine-induced neutralizing antibody and Fc-mediated effector responses share several qualitative features between rabbits and rhesus macaques, supporting the utility of rabbits for preclinical immunogen evaluation (55). In this study, rabbit immunizations demonstrated that the 330 Env immunogens elicited strong immune responses against autologous 330.16.E6 and the closely related pseudovirus 329.14.B1, consistent with the previous observation following 329 Env immunization (49). The high autologous neutralization may reflect the antigenic properties of 330.16.E6, including its intrinsic susceptibility to neutralization by multiple bnAbs and autologous plasma antibodies. However, the specific determinants responsible for the strong autologous response cannot be established from the present data (32). We compared the rabbit serum autologous neutralization titers elicited by the 330 Env with those reported for multiple HIV-1 SOSIP.664 immunogens, including 329 (49), ZM197 (11), IPGE-THIVC (24), AMC009 (12), B41 (13), AMC008 (13), BG505(v4.2) (13), and BG505(v5.2) (11) (**Table S9**). The 330 Env elicited comparatively higher levels of autologous neutralization than most of the evaluated Env trimers, except BG505, demonstrating its ability to induce potent strain-specific antibody responses in rabbits. However, only modest tier 1 heterologous neutralizing activity was observed, indicating limited breadth of the elicited response. High autologous neutralization titers can arise from antibodies targeting immunodominant, strain-specific surfaces or other highly accessible antigenic features, including glycan-deficient regions. They therefore should not be interpreted as evidence of progression toward bnAb responses. The absence of detectable cross-clade Tier-2 neutralization further supports the need for additional immunogen optimization to achieve broader neutralization. Consistent with previous immunization studies, the observed neutralization of MW965.26 and SF162 by 330 Env-elicited sera may suggest increased exposure of immunodominant epitopes, particularly within the V3 region (56). The multivalent presentation of 330 SOSIP.664 Env on ferritin NPs correlated with enhanced autologous neutralization compared with the soluble Env format. One possible explanation is increased B-cell receptor engagement resulting from repetitive antigen display (57); however, the underlying mechanism was not directly established in this study. Differences in antigen dose, particle trafficking, Env stability, antigen spacing or orientation, epitope accessibility, and avidity may also contribute to the observed response. Recently reported two-component NP platforms, including I53-50, dn2, and dn5, offer improved control over Env assembly and presentation compared with ferritin NPs and represent an important avenue for future investigation. Although the primary neutralization endpoint was Env-specific, inclusion of a ferritin-only control would have provided an additional assessment of scaffold-specific effects and helped determine whether anti-ferritin responses contributed to the observed binding responses. The absence of a ferritin-only control represents a limitation of the present study. The undetected cross-clade Tier-2 virus neutralization underscores the need to design further and optimize to broaden neutralization breadth. Further improvements in immunogenicity may be achieved through incorporation into in vitro self-assembling two-component nanoparticles, mRNA vaccine design approaches, evaluation of alternative adjuvants, and assessment of these vaccine candidates in non-human primate models. The immunization experiments were conducted exclusively in female rabbits with relatively small group sizes, which limits statistical power and the generalizability of the findings across sex and individual animals. Although rabbits are useful for evaluating HIV-1 Env immunogens, their B-cell repertoires, germline gene usage, and antibody maturation pathways differ from those in humans and may not fully reproduce the developmental pathways required for human glycan-dependent bnAb responses. Thus, the neutralization responses observed in rabbits provide evidence of immunogenicity but should not be interpreted as predictive of bnAb development in humans. Rabbit serum competition analysis suggested that immunization elicited polyclonal antibodies that competed with antibodies targeting several antigenic regions of the native-like 330 Env trimer, including the V1V2 apex, CD4bs, and V3-glycan supersite. However, these competition profiles do not establish that the elicited antibodies reproduce the V1V2-, CD4bs-, or V3-glycan-directed bnAb specificities observed in the AIIMS_330 donor (32). Strong competition against PGT145 and PGDM1400 suggests preservation of a well-ordered trimer apex, as these antibodies preferentially recognize quaternary V1V2 epitopes presented on properly folded Env trimers (58). The competition against VRC01 indicates the presence of antibodies targeting CD4bs, a key site of vulnerability frequently targeted during natural infection and by stabilized Env immunogens (17). Prominent competition against V3-glycan-directed bnAbs, including 44m, 10-1074, and PGT121, further supports presentation of glycan-dependent neutralizing determinants surrounding the N332 supersite and encouraged us to structurally characterize 330 Env in complex with autologous bnAb 44m. Although competition with non-nAbs was also detected, its magnitude was lower than that observed for bnAbs, consistent with previous Env immunization studies, in which both native and partially exposed non-neutralizing epitopes contribute to the overall antibody response (59). Definitive assignment of the elicited antibody specificities would require complementary approaches such as neutralization mapping with Env mutants, isolation and characterization of vaccine-elicited monoclonal antibodies, epitope-specific depletion experiments, or EM-based polyclonal epitope mapping (EMPEM)(60).

Structural characterization of HIV-1 Env trimers remains of great interest for structure-guided vaccine design. Given the limited number of high-resolution structures currently available for HIV-1 clade C Envs, including 16055 (51,52), DU172.17 (61), 426c (62), and CZA97.12 (63), the native-like 330 Env further expands the repertoire of structurally characterized clade C immunogens and provides additional structural insights into Env architecture, glycan organization, and antibody recognition. We observed the unliganded 330 Env adopted native-like structures with great structural similarity to clade A BG505 Env (RMSD gp120-0.685 Å & gp41-0.478 Å) than those previously reported for the clade B AMC009 Env (RMSD gp120-2.623 Å & gp41-1.624 Å), suggesting the ability of this Env to maintain prefusion closed conformations (12). The structural characterization of a contemporaneous pediatric EN-derived Env in complex with an autologous bnAb provides an uncommon opportunity to interrogate Env-antibody co-evolution within a naturally occurring broad-neutralization setting and to gain insight into the antigenic determinants of this Env, recognized by a co-evolving bnAb. Within the GDIR motif of the 330 Env, 44m interacted not only with the conserved D325 residue, but also with R327, whereas the previously solved BG505-44m structure showed predominant dependence on D325 alone (33). Further the structure suggests a possible interaction of the Env residues, E293, K337, and K446 with the bnAb 44m. To further understand the prevalence of the additional residues E293, K337, and K446 identified in this study, we aligned 18 HIV-1 Env variants generated from donor AIIMS_330 by single-genome amplification (SGA) in the previous study (32). This analysis revealed the conservation of key V3-glycan supersite elements, including the N332 glycan and GDIR motif, across all Env variants. In contrast, theese interface residues (E293, K337, and K446) were restricted to only four closely related Env variants E6, E1, E4, and E3 (**Figure S11**). The limited distribution of these residues within a subset of Envs suggests that accessory strain-specific residues may further shape the local epitope architecture required for optimal antibody engagement. Furthermore, the lack of neutralization dependendence on these residues, observed based on the neutralizing assys using mutant virsues, suggests that the precise role of these interacting residues, in terms of their antigenicity and immunogenicity, needs further evaluation. The precise role of these additional plausibly interacting residues in shaping immunogenicity remains to be investigated further.

Some of the limitations of this study are detailed as follows. are First, the relatively small group sizes of animals and the use of female rabbits alone limit the statistical power and generalizability of the immunogenicity findings. Second, the neutralization panel was limited and did not demonstrate detectable cross-clade Tier-2 breadth. Third, serum neutralization mapping was not performed, and therefore, the specific epitopes responsible for the observed neutralization responses remain unresolved. The glycan features of the Env immunogens were evaluated computationally rather than experimentally, limiting direct assessment of glycan occupancy and heterogeneity. In addition, PGT145-based purification may introduce selection for Env conformations preferentially recognized by this antibody. The absence of a ferritin-only control limits assessment of potential scaffold-specific contributions to the observed binding and immune responses. Finally, the 5.1 Å cryo-EM reconstruction limits confident assignment of individual side-chain contacts, hydrogen bonds, and glycan-mediated interactions. These limitations should be considered when interpreting the structural and immunogenicity findings and provide directions for further investigation.

In conclusion, our data establish that the 330 SOSIP.664 design adopts a native-like Env architecture and elicits a more potent autologous immunogenic response than the NFL.664 design. Multivalent presentation of 330 SOSIP.664 Env on ferritin was associated with a further increase in autologous neutralization, although the mechanism underlying this effect remains unresolved. The stabilized Indian clade C 330 Env may therefore serve as a useful antigenic probe for bnAb discovery and as a structural reagent for investigating Env–antibody recognition. Its incorporation into multivalent or sequential HIV-1 immunization strategies remains a future possibility that will require further optimization and validation, particularly to determine whether such approaches can improve neutralization breadth and elicit responses against Tier-2 viruses. Overall, the findings provide structural and antigenic insights into a pediatric elite-neutralizer-derived clade C Env and its recognition by the autologous 44m bnAb, which may inform future clade C HIV-1 immunogen design.

## MATERIALS AND METHODS

### Institutional Review Board & Ethics approval

This study was conducted after obtaining approval from the Institutional Ethics Committee of the All India Institute of Medical Sciences (AIIMS), New Delhi, India (250/IAEC-1/2020). Rabbit immunization was outsourced to a contract research organization (CRO), Liveon Biolabs Private Limited, Bengaluru, Karnataka, India, under an IAEC-approved protocol (No. LBPL-IAEC-008-01/2021; study number LBPL/NG-1736 (EF)). All animal experiments adhered to the guidelines for the care and use of laboratory animals as outlined by the Committee for Control and Supervision of Experiments on Animals (CCSEA), Government of India (64). The Env sequence and 44m antibody were originally obtained from an HIV-1 clade C-infected pediatric donor (AIIMS_330) after obtaining written informed consent from the parent/ guardian for collection. Approximately 5 mL of peripheral blood was collected from the donor. The original study for Env generation was approved by the Institute Ethics Committee of the All India Institute of Medical Sciences (AIIMS), New Delhi, under approval numbers IEC/NP-295/2011 and RP-15/2011. The subsequent antibody generation study was approved under IEC/59/08.01.2016, while the immunogen design study was approved under IECPG-354/29.05.2019.

### Construct design

The 330 Env gene was derived from a previously identified Indian clade C HIV-1 Env (330.16.E6, GenBank: MK076720) obtained from the pediatric elite neutralizer AIIMS_330, as previously described(32). The 330 Env construct was designed in soluble forms as SOSIP.664 (9) gp120-gp41-cleaved Env trimers and NFL.664 (65) gp140 single-chain Env trimers, with tailoring of SOSIP.664 Env trimers on self-assembling ferritin nanoparticles(26) to achieve multivalency. SOSIP.664 & NFL.664 constructs were stabilized by incorporating I559P & ΔMPER (Env prefusion stabilizing mutation)(66,67), I201C-A433C (intraprotomer locking disulfide bond, reduced V3 exposure)(10), and clade C stabilizing BG505-Trimer Derived (TD8) mutations (E47D, T49E, V65K, I165L, K429R, K432Q, K500R)(68). The SOSIP.664 construct was designed by introducing an SOS disulfide bond (A501C-T605C) (69), an I559P mutation (66), and an optimized furin cleavage site (RRRRRR)(70). In contrast, the NFL.664 (Native Flexible Linker) construct retained the I559P mutation and linked gp120 and gp41 using a flexible glycine-serine linker (GGGGSGGGGS)(65). The native signal sequence is replaced by the CD5 signal peptide (MGSLQPLATLYLLGMLVASVLA) for enhanced secretion of Envelope proteins. Constructs were codon-optimized for mammalian expression and synthesized commercially by GeneArt (Thermo Fisher Scientific). For binding assays, 330 Env constructs were tagged with either a His-tag (GSGSGGSGHHHHHHHH) or a twin strep tag (tst) (WSHPQFEKGGGSGGGSWSHPQFEK) at the C-terminus of gp41ECTO, preceding the stop codon. These constructs were subsequently cloned into the pPI4 expression vector using PstI and NotI restriction sites. The 330-ferritin NP construct was generated by fusing *Helicobacter pylori* ferritin (GenBank: WP_000949190.1), starting from Asp5 to Ser167, downstream of the SOSIP.664 C-terminus, separated by a Gly-Ser (GSG) linker, as described previously(26).

### Envelope Protein Expression

The 330 Env encoding plasmids were transiently expressed in Expi293F suspension cells (Gibco, A14527, RRID: CVCL_D615). Expi293F cells were thawed and maintained in Expi293 Expression Medium (Gibco, A1435101) in a shaker incubator (Eppendorf Brunswick, S41I230011) at 37°C, with 120 r.p.m. and 8% CO₂. For the production of SOSIP.664 Env, plasmids encoding SOSIP.664 Env (666.6 µg) and furin protease (333.3 µg) were mixed at a 3:1 ratio (w/w) for a 1-L Expi293F cell culture. The plasmid mixture was then incubated with PEI MAX transfection reagent (Polysciences, 24765) at a 3:1 (w/w) ratio of PEI MAX to total DNA for 20 min at room temperature and subsequently added to Expi293F cells at a density of 2 × 10⁶ cells/mL. For the production of the NFL.664 Env, NFL.664 plasmid DNA (1000 µg) was mixed with PEI MAX at a 3:1 ratio (w/w) and added to Expi293F cells under the same conditions. Seven days post-transfection, the culture supernatants were harvested, clarified by centrifugation, and filtered through 0.45-µm filters (Nalgene, 2954545).

### Envelope Protein Purification

330 Env proteins were purified by PGT145 bnAb immunoaffinity chromatography, as described previously (71). PGT145 affinity purification selectively enriches HIV-1 Env trimers with an intact quaternary apex epitope and was therefore used to isolate trimers in the prefusion, closed conformation. Briefly, HIV-1 pre-fusion closed trimer-specific bnAb PGT145 (72) was covalently coupled to the CNBr-activated Sepharose 4B beads (GE Healthcare, 17043001) according to the manufacturer’s protocol for antibody coupling. The PGT145-coupled CNBr Sepharose 4B bead column was used to purify HIV-1 Env trimers and ferritin NPs. Env proteins contained in Expi293F filtered supernatants were captured on PGT145-coupled CNBr-activated sepharose beads by overnight rolling incubation at 4 °C. Subsequently, the mixture of supernatants and beads was passed over Econo-columns (Bio-rad, 7371512). The columns were then washed with three column volumes of a 0.5 M NaCl and 20 mM Tris-HCl solution pH 8.0. After elution with 3 M MgCl_2,_ pH 7.5, the proteins were buffer-exchanged into TN75 (75 mM NaCl, 20 mM Tris-HCl, pH 8.0) by ultrafiltration using Vivaspin concentrators (MWCO 100 kDa; Cytiva, 28932363). Protein concentrations were determined from A280 values measured using a NanoDrop One (Thermo Fisher Scientific, 29018226), and molecular weight and extinction coefficient values were calculated by using ProtParam Expasy web tool. Env proteins used for immunization were first purified using PGT145 affinity chromatography, followed by size-exclusion chromatography (SEC) on a Superdex 200 Increase 10/300 GL column (GE Healthcare Life Sciences, 28990944) using an Äkta Pure 25M system (Cytiva, 29018226). Envs containing fractions were pooled, concentrated, filter-sterilized, snap-frozen in liquid nitrogen, and stored at -80°C until further use.

### SDS-PAGE and BN-PAGE Analyses

For SDS-PAGE and blue native polyacrylamide gel electrophoresis (BN-PAGE) analyses, 2 µg of 330 gp140 Env trimers, or equimolar amounts of ferritin NPs (2.5 µg), were run over a precast native gel 4–20 % Bis-Tris gradient gel (Bio-rad, 4561094). The degree of gp120-gp41 cleavage was examined by incubating the SOSIP.664 & NFL.664 proteins with 0.1 M dithiothreitol (DTT) and analyzed by SDS-PAGE under reducing conditions. Subsequently, the gels were stained with the Colloidal Blue Staining Kit (Invitrogen, LC6025).

### Enzyme-linked Immunosorbent assay (ELISA)

ELISAs were performed as previously described(73) using tst-tagged Env constructs on StrepTactinXT-coated microplates (IBA Lifesciences, 2-4101-001). Purified proteins (1 μg/mL) were diluted in NaHCO₃ buffer and immobilized on 96-well Streptactin plates by overnight incubation at 4°C. Following a three-wash step with PBST (0.05% Tween 20) to remove unbound trimers, plates were blocked with 1% BSA in PBST for 2 hours at RT. Further, the test antibodies were diluted in PBST/1% BSA and incubated for 2 h. After 3 washes with PBST, HRP-labeled goat anti-human IgG (Jackson ImmunoResearch, AB_2337578) diluted 1:3000 in PBST/1% BSA was added and incubated for 30 mins, followed by 3 washes with PBST and 3 washes with PBS. Ready-to-use 100μl of TMB substrate (Mabtech, 3652-F10) was then added to each well and incubated for 2 minutes at room temperature in the dark. The enzymatic reaction was stopped by adding 100μl of 2 N H₂SO₄ to each well. Absorbance was measured at 450 nm (OD450) using a 96-well microplate reader (Infinite 200 pro Tecan). Background absorbance from blank wells was subtracted from the corresponding sample readings. Binding curves were generated by plotting background-subtracted OD450 values against antibody concentration.

### Biolayer Interferometry (BLI)

Binding kinetics were measured using an Octet K2 (ForteBio, RED96e) device at room temperature with agitation at 1000 rpm. Anti-human Fc sensors (Sartorius AHC, 18-5060) were used to capture mAbs (10 μg/ml) for 300 sec, followed by equilibration in kinetics buffer for 60 s to obtain a baseline prior to analyte association. The 330 Envelope protein was used as an analyte in concentrations ranging from 500 nM to 6.1 nM in the HEPES buffer (pH 7.2) supplemented with 0.02% Tween 20 and 0.1% BSA. The mAb-captured sensors were immersed in wells containing the respective Env concentrations for 150 sec to measure association, followed by a 450 sec dissociation step in kinetics buffer. Blank wells without antibody loading were used as non-specific binding controls. The sensors were regenerated using three alternating washes of high pH (Kinetics buffer, pH 7.2) and low pH (1 M glycine, pH 2.4). Data were analyzed after subtracting from the reference sensor using the ForteBio Data analysis software (ForteBio Inc., Version 9.0). Sensorgrams were fitted using a 1:1 Langmuir binding model. Given the oligomeric nature of Env trimers and the multivalent display of Env on ferritin nanoparticles, the derived kinetic parameters are reported as apparent dissociation constants (*K*_D_^app^) and should be interpreted accordingly.

### Differential scanning fluorimetry (DSF)

330 SOSIP.664 protein was prepared at a concentration of 0.2 mg/mL in 1X PBS and transferred into 0.1 mL polypropylene PCR tube strips (Tarsons, 611020) with a final volume of 25 μL. Sypro Orange protein stain (Invitrogen, 4461146) was diluted 1:200. The samples were analyzed using a Rotor-Gene Q system (Qiagen, 9001862) with a high-resolution melt (HRM) protocol, starting at 25°C and increasing to 90°C in 1°C increments, with each step lasting 4 seconds. Melt curve data were processed using Rotor-Gene Q Series Software.

### Negative Stain Microscopy (ns-EM)

Negative-stain electron microscopy (ns-EM) was performed essentially as described elsewhere (74). Briefly, 3 μl of purified 330 gp140 Env trimers or ferritin NPs (□0.03 mg/mL) were applied to a glow-discharged (20 mA for 30 s) carbon-coated 400 Cu mesh grid. After 5 s, grids were negatively stained with 2% (w/v) uranyl formate for 60 s. Imaging was performed using a 120 kV Talos L120C electron microscope (TALOSL120CMS) equipped with a 4k x 4k Ceta camera at a magnification of 73kx (3.84 Å/pixel). The collected images were processed in EMAN 2.1(75). Particle extraction and reference-free 2D class averages were generated using simple_prime2D in SIMPLE 2.1 software (76) with a mask diameter of 30 pixels at 3.84 Å/pix.

### Immunizations

The rabbit immunization studies were conducted at a contract research organization (CRO), Liveon Biolabs Private Limited, Karnataka, India, under GLP-compliant conditions. Liveon Biolabs is an AAALAC International-accredited facility and is registered with the CCSEA, Government of India. Female, naive New Zealand White (NZW) rabbits, aged 4 months and weighing between 2.0 and 2.5 kg, were used for the study. The protocol for animal use was approved by the Institutional Animal Ethics Committee (IAEC) of Liveon Biolabs (Approval No.: LBPL-IAEC-008-01/2021; Study No.: LBPL/NG-1736 (EF)). Before housing, the experimental room was fumigated for decontamination, and the microbial load was assessed using the settle plate method. The rabbits were kept in a temperature-controlled Environment (20 ± 3 °C) with relative humidity ranging from 30–70%, a 12-hour light/dark cycle, 12–15 air changes per hour, and a noise level below 80 dBs. Mortality and morbidity were monitored twice daily, and clinical observations were conducted at least once daily throughout the study. Body weight measurements were recorded for all animals on day 1 before the administration of the test item and subsequently weekly (±1 day) during the treatment period. Female rabbits were randomly allocated into four groups. Group 1 immunized with SOSIP.664 Env gp140 trimers (n=4), Group 2 immunized with NFL Env gp140 trimers (n=4), Group 3 immunized with Ferritin Nanoparticles (n=4), and Group 4 served as Placebo (n=2), received Addavax adjuvant formulated with sterile 1X PBS. The gp140 SOSIP.664 and NFL.664 Env immunogens (G1 and G2) were administered at a dose of 20 µg of trimers, or an equivalent molar amount of SOSIP.664-ferritin nanoparticles (25 µg) (G3), formulated with the AddaVax adjuvant (1:1 v/v) (vac-adx-10, Invivogen). Doses were calculated based on the molecular weight of the protein peptides, excluding glycans, as described elsewhere (42). Briefly, the sequence-derived unglycosylated Env monomer and trimer molecular masses were 75.64 kDa and 226.91 kDa, respectively. The **H. pylori** ferritin monomer is ∼19.29 kDa, forming a 24-subunit scaffold of ∼462.9 kDa. Assuming the upper expected occupancy of 8 Env trimers per NP, the calculated molecular mass of the assembled NP is ∼2,278.1 kDa [462.9 + (8 × 226.91) kDa]. Thus, 20 µg of soluble Env trimer corresponds to 88.1 fmol of trimers, whereas 25 µg of ferritin nanoparticle corresponds to 11.0 fmol of NPs, equivalent to 87.8 fmol of Env trimers at 8 trimers per particle. Therefore, 25 µg of ferritin nanoparticle provides an approximately equimolar Env-trimer dose to 20 µg of soluble Env trimer. Rabbits were primed at week 0, and the first boost or second dose was administered at week 4, and the second boost or third dose was administered at week 20. Blood samples were collected from rabbits at weeks 0, 4, 6, 8, 12, 16, 20 & 22. Before blood collection from the marginal ear vein, a 4% lidocaine local anesthetic cream was applied for 5 minutes. The immunization process, including bleeding protocols, caused minimal, brief pain or distress and had no significant impact on animal health. Rabbits were euthanized by intravenous administration of an overdose of sodium thiopental (60 mg/kg) at the end of the immunization experiments, carried out by the CRO Liveon Biolabs (Approval No. LBPL-IAEC-008-01/2021; Study No. LBPL/NG-1736 (EF)). Complement-deactivated serum samples were subsequently analyzed for their binding reactivity and neutralization potential.

### HIV-1 pseudovirus generation

The HIV-1 pseudoviruses were produced in HEK293T cells (ATCC, CRL-3216; RRID: CVCL_0045) as described previously (77) by co-transfecting HIV-1 gp160 Envelope plasmid and a pSG3ΔEnv backbone plasmid. Briefly, 1 × 10^5^ cells in 2 mL complete DMEM (10% fetal bovine serum (FBS) (Gibco, 26140079 and 1% penicillin and streptomycin (Gibco, 15140-122) were seeded per well of a 6-well cell culture plate (Costar, 3506) the day before transfection. For transfection, Envelope (1.25 μg) and delta Envelope (pSG3ΔEnv backbone) plasmids (2.50 μg) were mixed in a 1:2 ratio in Opti-MEM (Gibco, 31985062), with a final volume of 200 μl per well, and incubated for 5 min at room temperature. Next, 3 μl of PEI-Max transfection reagent (Polysciences, 24765) (1 mg/mL) was added to this mixture before further incubation for 15 min at room temperature. This mixture was then added dropwise to HEK 293T cells supplemented with fresh complete DMEM, and the cells were incubated at 37 °C for 48 h. Pseudoviruses were then harvested by filtering cell supernatants with 0.45 μm sterile filters (MDI), aliquoted, and stored at −80 °C until usage.

### HIV-1 neutralization assay

Neutralization assays were carried out using TZM-bl reporter cells, as described elsewhere (78). The autologous and heterologous antibody responses in endpoint sera were measured in TZM-bl cell neutralization assays. Neutralization was measured as a reduction in luciferase gene expression after a single round of infection of TZM-bl cells (NIH AIDS Reagent Program, 8129; RRID: CVCL_B478) with HIV-1 Env pseudoviruses. The TCID_50_ of the HIV-1 pseudoviruses was calculated, and 200 TCID_50_ units were used in the neutralization assays by incubation with three-fold serially diluted rabbit sera starting at a 1:20 dilution. Next, freshly trypsinized TZM-bl cells in a complete DMEM (10% fetal bovine serum (FBS) (Gibco, 26140079 and 1% penicillin and streptomycin (Gibco, 15140-122) containing 50 μg/mL DEAE-dextran (Sigma, D9885) at 10^5^ cells/well were added, and the plates were incubated at 37 °C for 48 h. Virus controls (cells with HIV-1 virus only) and cell controls (cells without virus and antibody) were included. MuLV was used as a negative control. After 48 hours of incubation, luciferase activity was measured using the Bright-Glow Luciferase Assay System (Promega, E2610). ID_50_ for antibodies was calculated from a dose–response curve fit using a non-linear function using GraphPad Prism 8 (San Diego, CA, USA). All neutralization assays were repeated at least 2 times, and ID_50_ values were calculated independently for each assay replicate. The replicate ID_50_ values were averaged and reported as the mean ID_50_ in Supplementary **Table S3**.

### Site-Directed Mutagenesis

Site-directed mutagenesis (SDM) was performed to assess the contribution of Env residues identified at the 44m bnAb–330 Env interface by cryo-EM structural analysis. Alanine-substitution mutants were generated for the selected residues (E293, D325, R327, H330, N332, K337, and K446) using the Agilent QuikChange site-directed mutagenesis protocol/software. Mutagenic primers were designed using the manufacturer’s recommended parameters and synthesized commercially. PCR amplification was performed using 330.16.E6 as the template. The cycling conditions consisted of an initial denaturation at 94°C for 1 min, followed by denaturation at 94°C for 30 s, primer annealing at 30 s, and extension at 72 °C for 4.5 min, with a final extension at 72°C for 9 min. Following amplification, PCR products were analyzed by agarose gel electrophoresis and treated with 1 U of DpnI to digest the parental methylated template DNA. The resulting products were transformed into chemically competent STBL3 **E. coli** cells. Individual colonies were selected, and plasmid DNA was isolated and subjected to commercial Sanger sequencing to confirm the presence of the desired mutations. Sequence confirmed Env mutants were transfected into HEK293T cells for pseudovirus production. The resulting pseudoviruses were evaluated for neutralization sensitivity using the TZM-bl-based neutralization assay as described above.

### Serum competition assay

Serum competition analysis was performed using biolayer interferometry (BLI)-based tandem epitope binning based on the Octet platform (ForteBio, RED96e). Briefly, the serum samples from individual animals were pooled by the SOSIP.664 and NFl.664 groups and diluted 1:3 to a final volume of 200 µL per well for serum pre-saturation of the Env antigen. The twin-strep-tagged HIV-1 Env antigens (SOSIP.664 and NFL.664 trimers) were immobilized onto High Precision Streptavidin (SAX) biosensors (Sartorius, 18-5117) at 10 µg/mL for 30s, followed by a 30s baseline step in kinetics buffer. Biosensor-bound antigens were first saturated with polyclonal immune sera for 330 s and subsequently subjected for competition to HIV-1 broadly neutralizing, including V1V2 apex-directed (PGT145 and PGDM1400), CD4bs-directed (VRC01), V3-glycan supersite-directed (44m, 10-1074, and PGT121), and the gp120-gp41 interface-directed (PGT151), along with non-neutralizing antibodies (F105, 447-52D, 412d, and 17b) for 100 s to assess residual binding. Percent inhibition was calculated using the formula[1 — (R(serum + mAb) / R(mAb only))] \times 100. Rserum + mAb represents the residual binding response of the monoclonal antibody following serum pre-saturation of the Env antigen and RmAb only represents monoclonal antibody binding in the absence of serum. A threshold of 35% inhibition was applied to minimize the influence of background variation and facilitate interpretation of the serum competition matrix, which was analyzed using ForteBio Data analysis software (ForteBio Inc., Version 9.0). Percent inhibition was calculated using the formula [1 — (R(serum + mAb) / R(mAb only))] \times 100. The calculated inhibition values are globally normalized to a 0–10 scale using min–max normalization, with the minimum (−9.7) and maximum (215.9) values across all mAbs and immunogen combinations assigned scores of 0 and 10, respectively. Intermediate responses were proportionally scaled between these values and visualized as a heatmap.

### Preparation of 44m Fab fragment

Fab fragments were generated from 4 mg of AIIMS-44m IgG using a Fab fragmentation kit (G Biosciences, 786-273) following the manufacturer’s protocol. Briefly, 0.5 ml of 44m IgG was desalted using a SpinOUT GT-600 column. Cysteine.HCl was added to the Fab Digestion Buffer (pH=7) to a final concentration of 20 mM before papain digestion. The 0.5 ml IgG sample was then incubated with Immobilized Papain in the spin column at 37°C for 6 hours with end-over-end mixing. After incubation, the digested antibody solution was collected by centrifugation at 5,000 ×g for 1 minute. The flow-through was further incubated with Protein-A resin for 10-15 minutes at room temperature (RT) with end-over-end mixing. Subsequently, the Fab fragments were collected in the flow-through while Fc fragments and undigested IgG remained bound to the resin. The purity and size of the Fab fragments were assessed by SDS-PAGE, and the purified Fab fragments were stored at -80°C.

### Cryo-EM sample preparation, data acquisition, data analysis, and model fitting

The 330 Env trimer (1.5 mg/ml) was incubated with 44m Fab at a 1:3 molar ratio. A 3 μl sample was applied to glow-discharged R1.2/R1.3 300-mesh copper grids and immediately blotted for 6 seconds with a blotting force of 5 using the FEI Vitrobot Mark IV plunger at 22°C and 100% humidity. The grid was immediately plunge-frozen into liquid ethane at liquid nitrogen temperature. Cryo-EM data were collected using a 200□kV Talos Arctica transmission electron microscope (Thermo Scientific, TALOSARCTICA) equipped with a Gatan K2 Summit Direct Electron Detector. Movies were recorded automatically using Latitude-S (DigitalMicrograph - GMS 3.5) at a nominal magnification of 45,000x at the effective pixel size of 1.17□Å (76). Micrographs were acquired in counting mode with a total dose of 40 e^-^/Å^2^, and an exposure time of 8 sec, distributed across 20 frames. A total of 6046 movies were acquired for the 330 SOSIP.664 Env trimer-44m Fab complex datasets. Single-Particle Analysis (SPA) was performed on the acquired Cryo-EM movies using cryoSPARC 4.5 (79). □ At first, drift and gain corrections for the individual movies were applied using Patch Motion Correction, and Contrast Transfer Function (CTF) parameters were estimated using Patch CTF estimation.□Subsequently, CTF-estimated micrographs were analyzed to eliminate low-resolution micrographs, using a fit resolution threshold of 8 Å and curated exposure. The particles from the best micrographs were selected for automated picking using a 2D reference in Relion and extracted with a box size of 320□pixels for the 44m Fab-bound 330 SOSIP.664 Env trimer complexes. The extracted particles were imported into cryoSPARC v4.5, and the remaining processing was performed in that software. After four rounds of rigorous 2D classification, 363,761 particles were selected from the best 2D classes with high-resolution features of the Ab-bound complex. This dataset was divided into 4 classes for Ab initio reconstruction with C3 symmetry. Class 04, with 118572 particles, had the best Ab-bound features among all the classes (**Figure S6**). Therefore, we considered class 04 for non-uniform refinement by imposing C3 symmetry. The resolution of the 3D map was estimated at FSC 0.143 Å. The local-resolution estimation tool in cryoSPARC was used to estimate local resolution. The AlphaFold model of the AIIMS 330 env and the 44m bnAb were fitted to the cryo-EM map in ChimeraX (80) and Coot (81). The AlphaFold model of the AIIMS 330 env and the 44m bnAb were used to build atomic models using Phenix 1.19.2 (82). For model fitting, the Cryo-EM maps of 330 Env trimers were docked onto the ferritin structure (PDB: 3BVE) at four symmetric sites. Overviews of Cryo-EM data processing are shown in Table S5. The atomic model fitting was done and visualized using ChimeraX (80). We used the PDBsum generate server (83) to perform interface area calculations, and ChimeraX (80) for electrostatic analysis, contact assignment, and RMSD analysis. The same data collection and processing parameters were used for 330 Env trimers (Supplementary table S8).

### Statistical analysis

Statistical analyses were performed using GraphPad Prism v9.5.1 (GraphPad Software, La Jolla, CA, USA). Differences among the SOSIP.664, NFL.664, and Ferritin NP groups were evaluated using unpaired two-tailed *t*-tests or one-way ANOVA followed by Tukey’s multiple-comparisons test, as appropriate. Longitudinal serum-binding responses were analyzed using linear mixed-effects models (LMMs) to account for repeated measurements within animals and to assess the effects of immunogen group, time, and group × time interaction. Statistical significance was defined as *P* < 0.05, and exact *P* values are reported where applicable. Data are presented as mean ± standard deviation (SD). Neutralization titers are presented as median with interquartile range (IQR). Experiments were performed in duplicates and independently repeated twice.

## Supporting information

Supplementary Figures and Tables version 3

## ACKNOWLEDGMENTS

The authors express gratitude to Prof. Rogier W. Sanders for generously providing the pPI4 expression vector and for training S.D.S. as part of his HIV-1 Trust Fellowship. We are grateful to the NIH AIDS reagent program for providing the HIV-1 research reagents and the Neutralizing Antibody Consortium (NAC), IAVI, USA, for providing the HIV-1 neutralizing antibodies donated by Michel Nussenzweig, Hermann Katinger, Mark Connors, James Robinson, Dennis Burton, John R. Mascola, Peter D. Kwong, and William Olson. We appreciate Liveon Biolabs’ support in conducting the immunization studies. We also extend our appreciation for the IAVI-Leader Development Program awarded to S.D.S.

## AUTHOR CONTRIBUTIONS

**Swarandeep Singh:** Conceptualization, Methodology, Validation, Formal analysis, Data Curation, Investigation, Writing - Original Draft, Visualization, Project administration **Sanjeev Kumar:** Conceptualization, Validation, Formal analysis, Writing - Review & Editing, Visualization, Project administration, Funding acquisition **Arnab Chatterjee**: Methodology, Validation, Formal analysis, Data Curation, Investigation, Writing - Original Draft, Project administration **Lakshay Malhotra:** Methodology, Validation, Investigation, Data Curation **Muneeb Pervez:** Methodology, Validation, Investigation, Data Curation **Abdul Rahman:** Methodology, Validation, Investigation, Data Curation **Ifrah Andrabi:** Methodology, Validation, Investigation, Data Curation **Bapan Mondal:** Methodology, Validation, Investigation, Data Curation **Sanket Katpara:** Methodology, Validation, Investigation, Data Curation **Shaifali Sharma:** Methodology, Validation, Investigation, Data Curation **Harsh Bhakhri:** Methodology, Validation, Investigation, Data Curation **Anjli Gaur:** Methodology, Validation, Investigation, Data Curation **Tanu Bansal:** Methodology, Validation, Investigation, Data Curation **Vikram Iyer:** Methodology, Validation, Investigation, Data Curation **Abdul Wahid Hussain:** Methodology, Validation, Investigation, Data Curation **Shubbir Ahmed:** Methodology, Validation, Supervision, Writing - Review & Editing **Rajesh Kumar:** Methodology, Validation, Supervision, Writing - Review & Editing **Ethayathulla Abdul Samath:** Methodology, Validation, Supervision, Writing - Review & Editing **Rakesh Lodha:** Supervision, Resources, Writing - Review & Editing **Somnath Dutta:** Methodology, Validation, Formal analysis, Resources, Writing - Review & Editing, Funding acquisition **Kalpana Luthra:** Conceptualization, Validation, Resources, Writing - Review & Editing, Supervision, Funding acquisition

## DATA AVAILABILITY

The modified 330 Envelope protomer sequences have been deposited in GenBank under accession numbers: PV575182 (330_SOSIP.664), PV575183 (330_NFL.664), and PV575184 330_SOSIP.664 ferritin Nanoparticle). The cryo-EM maps and atomic coordinates of the 330 SOSIP.664 Env trimer in complex with bnAb 44m has been deposited in the Electron Microscopy Data Bank (EMDB), and the Protein Data Bank (PDB) under accession codes EMD-66717 and 9XC1, respectively. Additional data are available from the corresponding authors upon reasonable request.

## ABBREVIATIONS

AUC: Area Under Curve
BLI: Biolayer Interferometry
bnAb: Broadly Neutralizing Antibodies
CD4bs: CD4 Binding Site
DSF: Differential Scanning Fluorimetry
ELISA: Enzyme Linked Immuno Sorbent Assay
EM: Electron Microscopy
EMPEM: Electron Microscopy-based Polyclonal Epitope Mapping
EN: Elite Neutralizer
Env: Envelope
NFL: Native Flexible Linker
nAb: Neutralizing Antibodies
non-nAbs: Non-Neutralizing Antibodies
NP: Nanoparticle
NS: Negative Stain
PNGS: Potential N-Linked Glycosylation Sites
RMSD: Root Mean Square Deviation
SEC: Size Exclusion Chromatography
TMB: 3,3□,5,5□-Tetramethylbenzidine

