## Supplementary Figures and Tables version 3 for "Immunogenicity and Structure of a Stabilized HIV-1 clade C Env from a Pediatric Elite Neutralizer in Complex with an Autologous bnAb"

```
                ....|....| ....|....| ....|....| ....|....| ....|....| ....|....| ....|....| ....|....|
                        10         20         30         40         50         60         70         80
HXB2            MRVKEKYQHL WRWGWRWGTM LLGMLMICSA TEKLWVTVYY GVPVWKEATT TLFCASDAKA YDTEVHNVWA THACVPTDPN
330.16.E6       MKVRGTLRNY RQW-WIWGVL GFWMLMICNV EGDSWVTVYY GVPVWKEAKT TLFCASDAKA YDKEVHNVWA THACVPTDPS
330.NFL         ---MGSLQPL ATL-YLLGML ----VASVLA AGNLWVTVYY GVPVWKDAET TLFCASDAKA YDKEKHNVWA THACVPTDPS
330.SOSIP       ---MGSLQPL ATL-YLLGML ----VASVLA AGNLWVTVYY GVPVWKDAET TLFCASDAKA YDKEKHNVWA THACVPTDPS
Consensus            . :        :  * :      :    .    . ****** ******:* * ********** **.* ***** *********.

                ....|....| ....|....| ....|....| ....|....| ....|....| ....|....| ....|....| ....|....|
                        90        100        110        120        130        140        150        160
HXB2            PQEVVLVNVT ENFNMWKNDM VEQMHEDIIS LWDQSLKPCV KLTPLCVSLK CTDLKNDTNT NSSSGRMIME KGEIKNCSFN
330.16.E6       PQEMVLENVT ENFNMWKNDM VDQMQEDVIS LWDQSLKPCV KLTPLCVTLI CANVNSNSST TLN---ETHD GSELKNCSFN
330.NFL         PQEMVLENVT ENFNMWKNDM VDQMQTDVIS LWDQSLKPCV KLTPLCVTLI CANVNSNSST TLN---ETHD GSELKNCSFN
330.SOSIP       PQEMVLENVT ENFNMWKNDM VDQMQTDVIS LWDQSLKPCV KLIPLWVTLI CANGNSKSST TLN---ETHD GSELKNCSFN
Consensus       ***:** *** ********** *:**: *:** ********** ** ** *:*   *:: :..:.* . .      :  .*:******

                ....|....| ....|....| ....|....| ....|....| ....|....| ....|....| ....|....| ....|....|
                       170        180        190        200        210        220        230        240
HXB2            ISTSIRGKVQ KEYAFFYKLD IIPIDN---D TTSYKLTSCN TSVITQACPK VSFEPIPIHY CAPAGFAILK CNNKTFNGTG
330.16.E6       ATTEIRDRKQ TVHALFYRLD ILPLDEKNKN SSKYRLINCN TSAITQACPK VTFDPIPIHY CAPAGYAILK CNNKTFNGTG
330.NFL         ATTELRDRKQ KVHALFYRLD ILPLDEKNKN SSKYRLINCN TSACTQACPK VTFDPIPIHY CAPAGYAILK CNNKTFNGTG
330.SOSIP       GTTELRDRKQ KVHALFYRLD ILPLDEKNKN SSKYRLINCN TSACTQACPK VTFDPIPIHY CAPAGYAILK CNNKTFNGTG
Consensus        :*.:*.: * . :*:**:** *:*:*:    : ::.*:* .** **. ****** *:*:****** *****:**** **********

                ....|....| ....|....| ....|....| ....|....| ....|....| ....|....| ....|....| ....|....|
                       250        260        270        280        290        300        310        320
HXB2            PCTNVSTVQC THGIRPVVST QLLLNGSLAE EEVVIRSVNF TDNAKTIIVQ LNTSVEINCT RPNNNTRKRI RIQRGPGRAF
330.16.E6       PCHNVSTVQC THGIKPVVST QLLLNGSLAE GEIIIRSENL TNNVKTIIVH LNQSVEIVCT RPNNNTRKSI RIGPGQ-TFY
330.NFL         PCHNVSTVQC THGIKPVVST QLLLNGSLAE GEIIIRSENL TNNVKTIIVH LNQSVEIVCT RPNNYTRKSI RIGPGQ-TFY
330.SOSIP       PCHNVSTVQC THGIKPVVST QLLLNGSLAE GEIIIRSENL TNNVKTIIVH LNQSVEIVCT RPNNYTRKSI RIGPGQ-TFY
Consensus       ** ******* ****:***** **********  *::*** *: *:*.*****: ** **** ** **** *** * **  *    :

                ....|....| ....|....| ....|....| ....|....| ....|....| ....|....| ....|....| ....|....|
                       330        340        350        360        370        380        390        400
HXB2            VTIGKIGNMR QAHCNISRAK WNNTLKQIAS KLREQFGNNK TIIFKQSSGG DPEIVTHSFN CGGEFFYCNS TQLFNSTWFN
330.16.E6       AT-DIIGDIR QAHCNISEKK WHETLKGVGK KLAEHFPN-K TIKFTSSSGG DLEITMHSFN CRGEFFYCNT SGLFNSTYNG
330.NFL         ATGDIIGDIR QAHCNISEKK WHETLKGVGK KLAEHFPN-K TIKFTSSSGG DLEITMHSFN CRGEFFYCNT SGLFNSTYNG
330.SOSIP       ATGDIIGDIR QAHCNISEKK WHETLKGVGK KLAEHFPN-K TIKFTSSSGG DLEITMHSFN CRGEFFYCNT SGLFNSTYNG
Consensus       .* . **::* *******. * *::*** :..  ** *:* * * ** *..**** * **. **** * *******: : *****: .

                ....|....| ....|....| ....|....| ....|....| ....|....| ....|....| ....|....| ....|....|
                       410        420        430        440        450        460        470        480
HXB2            STWSTEGSNN TEGS-----D TITLPCRIKQ IINMWQKVGK AMYAPPISGQ IRCSSNITGL LLTRDGGNSN --NESEIFRP
330.16.E6       TSGLFNSTYN GTYSHDGVNS TITLPCRIKQ IINMWQEVGR AMYAPPIEGN ITCKSNITGI LLVRDGGDNN -QSDTETFRP
330.NFL         TSGLFNSTYN ---------S TITLPCRIKQ IINMWQRVGQ CMYAPPIEGN ITCKSNITGI LLVRDGGDNN -QSDTETFRP
330.SOSIP       TSGLFNSTYN ---------S TITLPCRIKQ IINMWQRVGQ CMYAPPIEGN ITCKSNITGI LLVRDGGDNN HQSDTETFRP
Consensus       ::   :.: *          . ********** ******.**: .******.*: * *.*****: **.****:.*    .::* ***

                ....|....| ....|....| ....|....| ....|....| ....|....| ....|....| ....|....| ....|....|
                       490        500        510        520        530        540        550        560
HXB2            GGGDMRDNWR SELYKYKVVK IEPLGVAPTK AKRRVVQ--- ---REKRAVG IGALFLGFLG AAGSTMGAAS MTLTVQARQL
330.16.E6       GGGDMRDNWR SELYKYKVVE IKPLGVAPTA AKRRVVE--- ---RGKRAVG IGAVFLGFLG AAGSTMGAAS ITLTVQARQL
330.NFL         GGGDMRDNWR SELYKYKVVE IKPLGVAPTR AKRRVVEGGG GSGGGGSAVG IGAVRRGFLG AAGSTMGAAS ITLTVQARQL
330.SOSIP       GGGDMRDNWR SELYKYKVVE IKPLGVAPTR CKRRVVE--- -RRRRRRAVG IGAVSLGFLG AAGSTMGAAS MTLTVQARNL
Consensus       ********** *********: *:*******  .*****:            *** ***:  **** ********** :*******:*

                ....|....| ....|....| ....|....| ....|....| ....|....| ....|....| ....|....| ....|....|
                       570        580        590        600        610        620        630        640
HXB2            LSGIVQQQNN LLRAIEAQQH LLQLTVWGIK QLQARILAVE RYLKDQQLLG IWGCSGKLIC TTAVPWNASW SNKSLEQIWN
330.16.E6       LSGIVQQQSN LLRAIEAQQH LLQLTVWGIK QLQTRVLAIE RYLKDQQLLG LWGCSGKLIC TTAVPWNSSW SNKSQEDIWN
330.NFL         LSGIVQQQSN LPRAPEAQQH LLQLGVWGIK QLQTRVLAIE RYLKDQQLLG LWGCSGKLIC TTAVPWNSSW SNKSQEDIWN
330.SOSIP       LSGIVQQQSN LLRAPECQQH LLQDTHWGIK QLQTRVLAIE HYLKDQQLLG LWGCSGKLIC CTAVPWNSSW SNKSQEDIWN
Consensus       ********.* * ** *.*** ***   **** ***:*:**:* :********* :*********  ******:** **** *:***

                ....|....| ....|....| ....|....| ....|....|
                       650        660        670        680
HXB2            HTTWMEWDRE INNYTSLIHS LIEESQNQQE KNEQELLELD
330.16.E6       NMTWMQWDRE INNYTNTIYQ LLEESQNQQE KNEKDLLALD
330.NFL         NMTWMQWDRE IGNYTNTIYQ LLEESQNQQE KNEVDLLALD
330.SOSIP       NMTWMQWDRE IGNYTNTIYQ LLEESQNQQE KNEVDLLALD
Consensus       : ***:**** *.***. *:. *:******** *** :** **
```

**Figure S1. Sequence alignment of 330 gp140 SOSIP and NFL amino acid sequences to the HXB2 reference coordinates.**

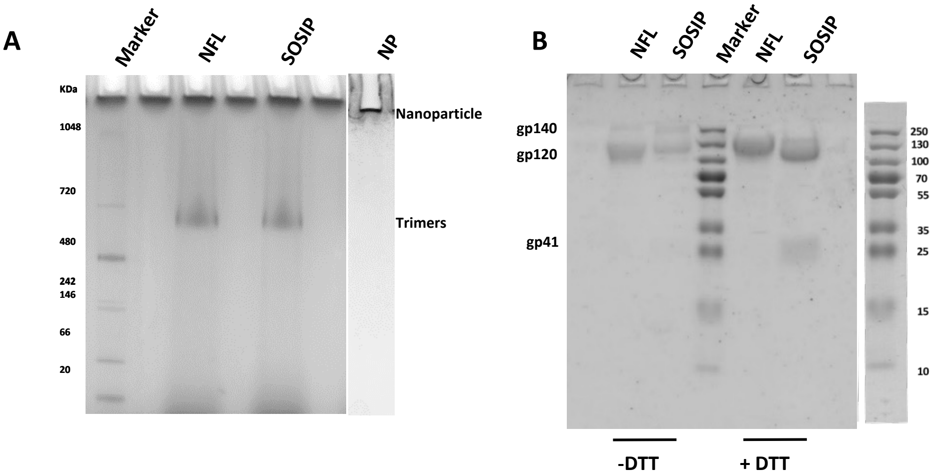

**Figure S2. Biochemical characterization of 330 gp140 Env trimers and ferritin nanoparticles. (A)** Blue Native-PAGE analysis of PGT145 purified 330 gp140 SOSIP and NFL trimers, along with 330 ferritin NPs, stained with Coomassie blue. The soluble Env trimers migrated as discrete high-molecular-weight bands consistent with intact trimeric species, whereas the ferritin NPs remained predominantly at the top of the gel because of their large molecular size. **(B)** SDS-PAGE analysis of purified 330 gp140 Env trimers under reducing and non-reducing conditions. Under reducing conditions (+DTT), the Env proteins migrated predominantly as monomeric gp120 and gp41 species, indicating the purity and integrity of the trimer preparations.

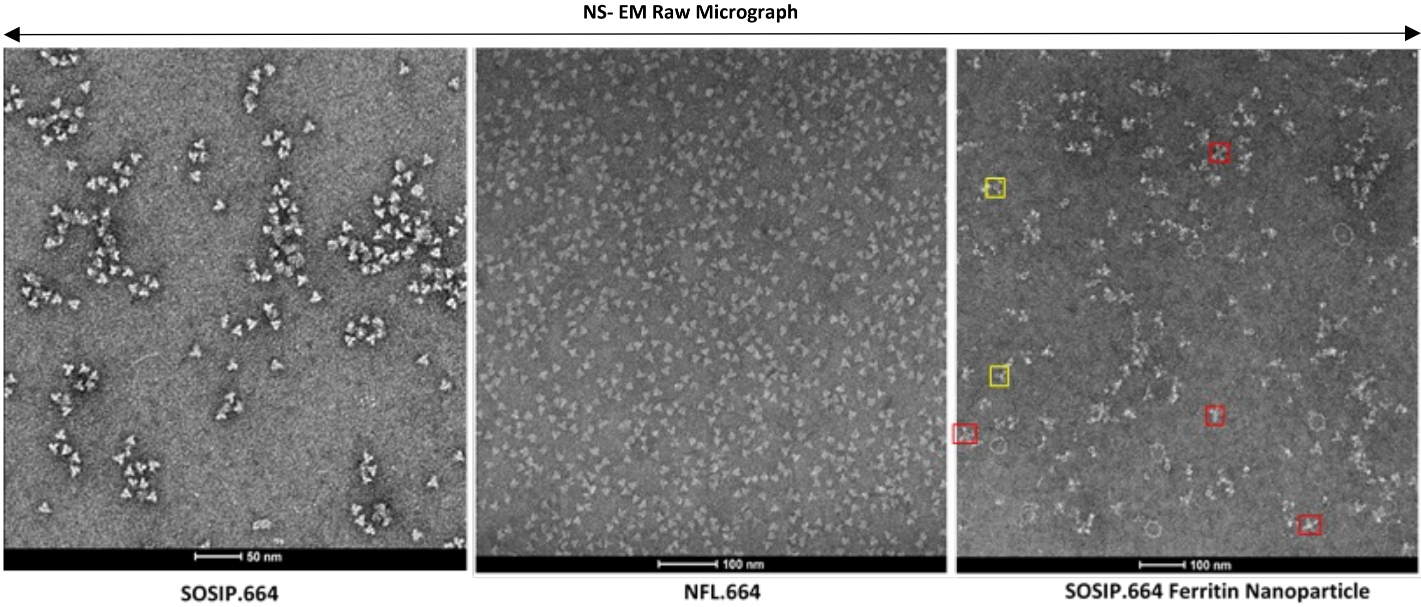

**Figure S3. Negative staining electron microscopy raw micrographs. (A)** Representative negative-stain TEM (nsTEM) raw micrographs of 330 SOSIP env trimers at 50 nm scale. **(B)** Raw micrograph representing 330 NFL env trimers at 100 nm scale. **(C)** Representative micrographs of self-assembling H. pylori ferritin nanoparticles presenting the SOSIP Env trimers, red indicating four or more trimers presentation, while yellow indicates up to two Env presentations by ferritin particles.

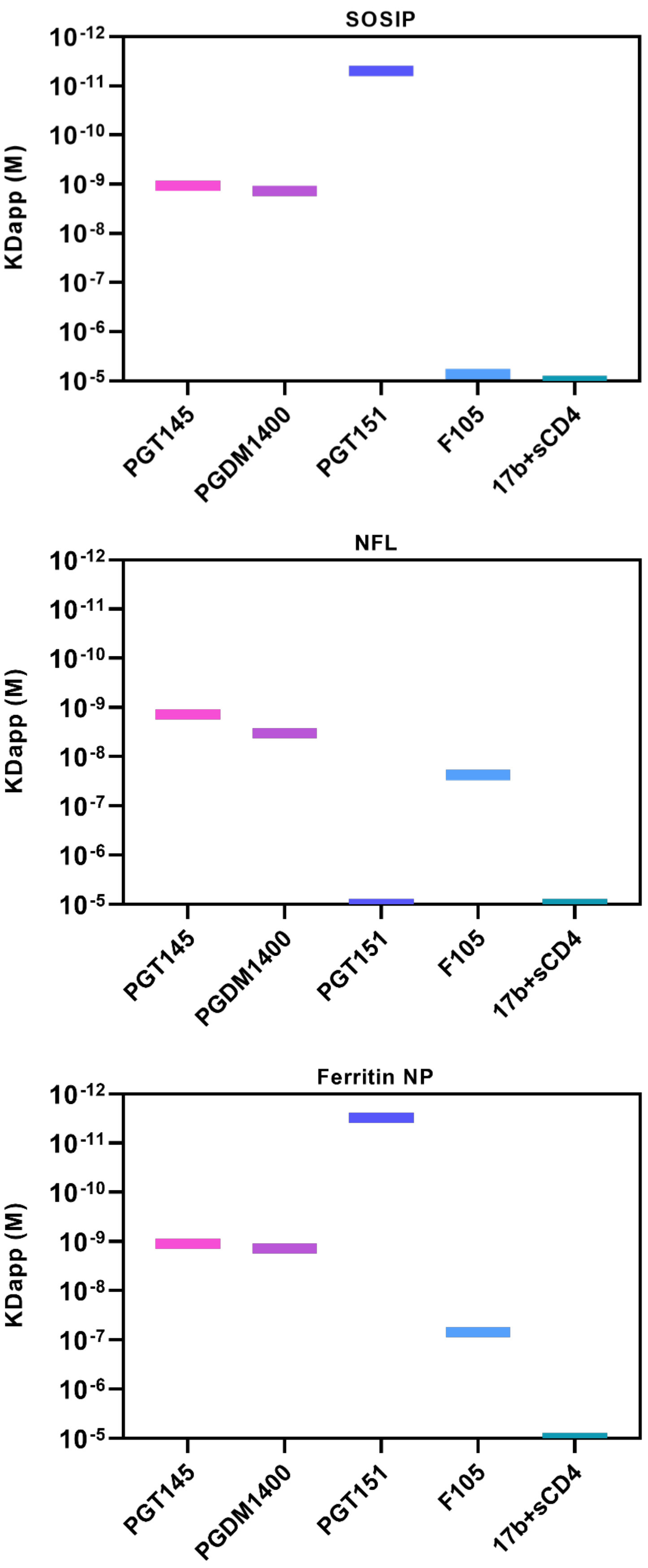

**Figure S4. BLI-Octet analysis of antibody binding to 330 gp140 Env trimers and ferritin nanoparticles.** Binding affinities of 330 SOSIP, NFL, and ferritin NPs to bnAbs PGT145, PGDM1400, and PGT151 and non-nAbs F105 and 17b in the presence of sCD4 were determined by biolayer interferometry (BLI). Apparent $K_D$ values ($K_D^{app}$ ) are shown on a logarithmic scale in molar (M) units. The KD values were derived using 1:1 global fitting.

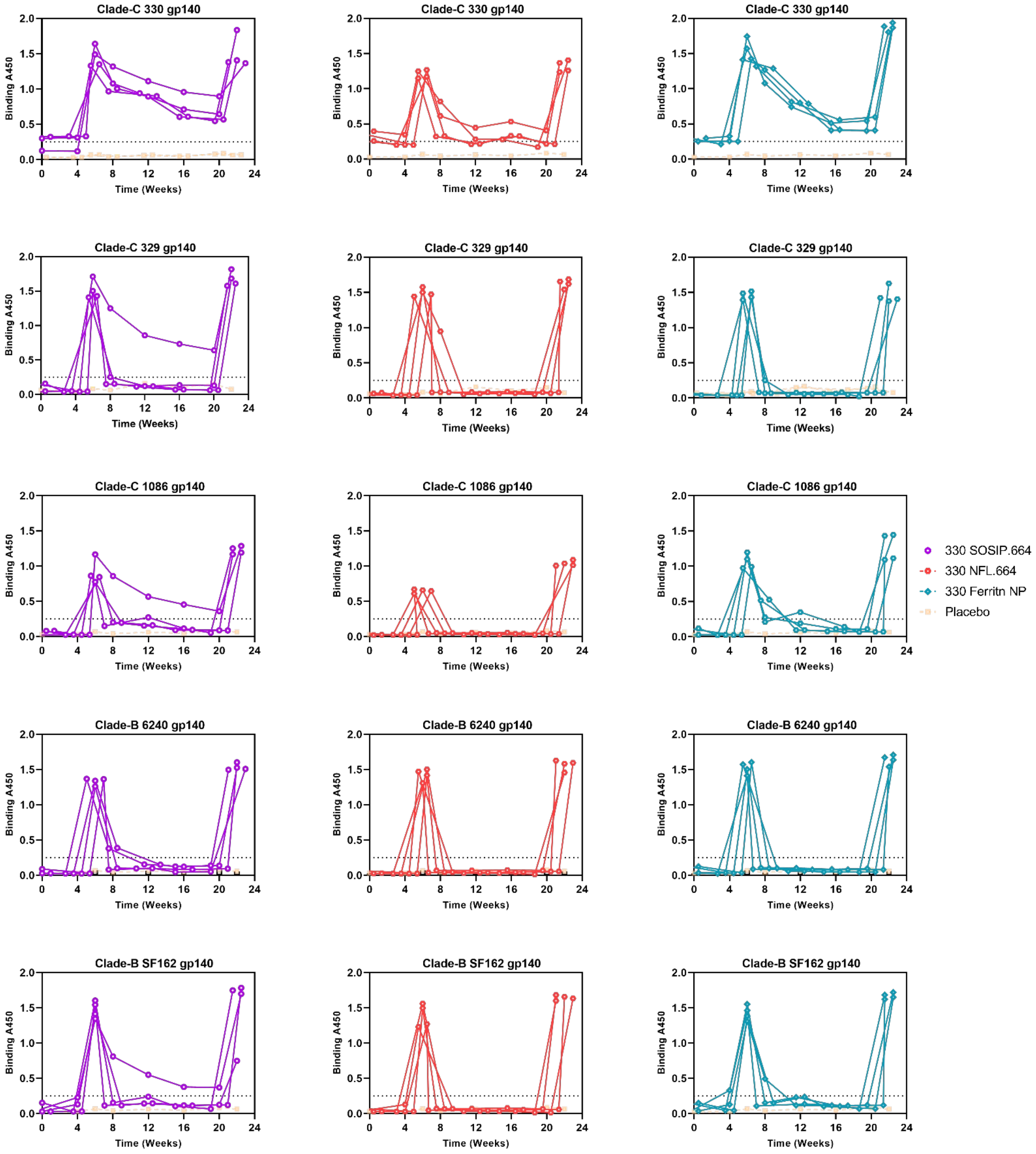

**Figure S5. Longitudinal binding endpoint titers of individual animals in each group.** Serum antibody endpoint titers were determined longitudinally by ELISA against the clade C autologous Env 330 and heterologous HIV-1 gp140 Env proteins, including clade C Envs 329 and 1086 and clade B Envs SF162 and 6240. Individual animal endpoint titers are shown for each time point.

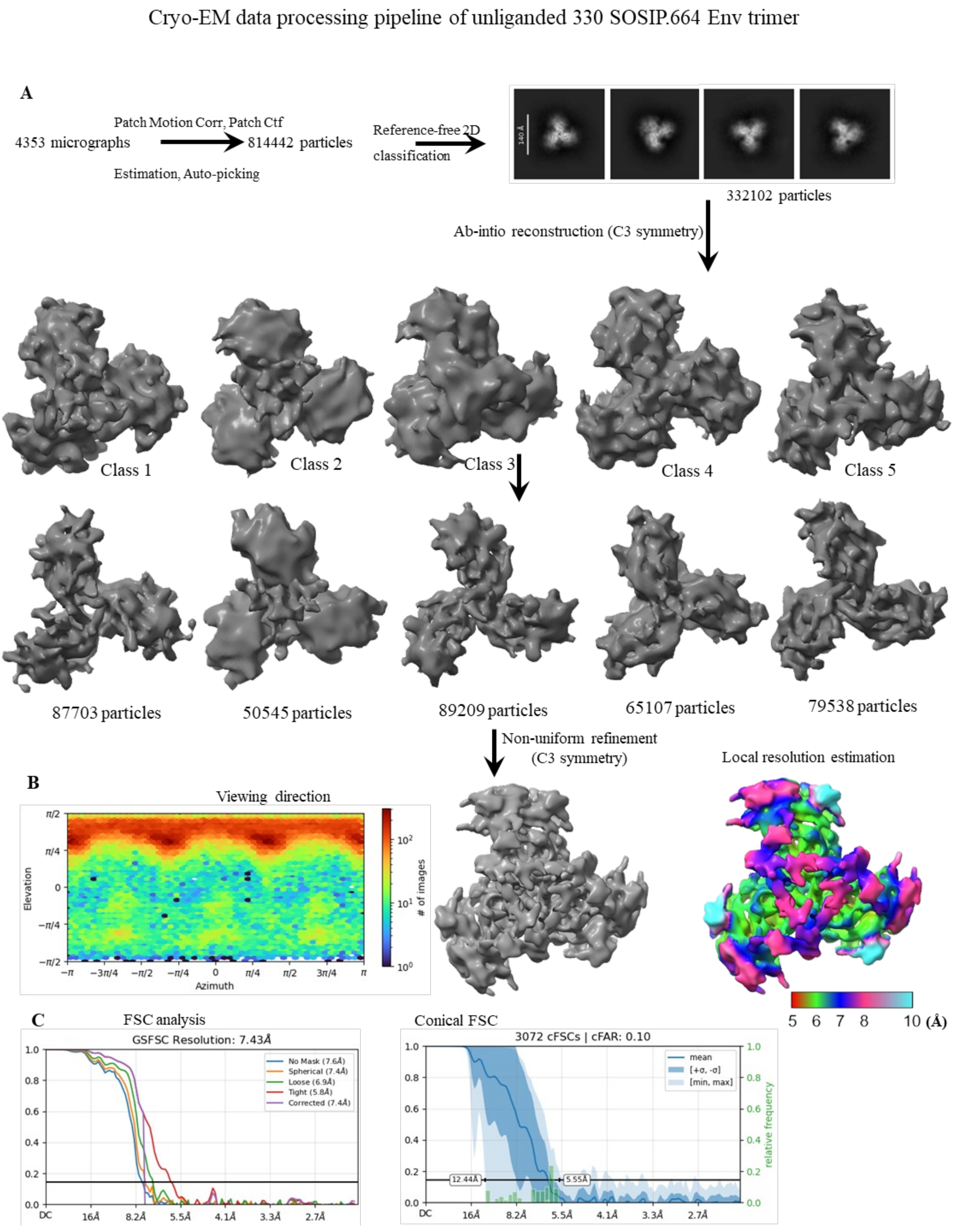

**Figure S6: Cryo-EM analysis of unliganded 330 SOSIP.664 Env trimer (A)** Pipeline of Cryo-EM data processing using single particle cryo-EM and 3D classification of 330 SOSIP.664 Env trimer. **(B)** The viewing-direction particle distribution for Class 04 reveals diverse particle orientations in the dataset. **(C)** Gold standard Fourier Shell Correlation (FSC) curve of the cryo-EM map at 7.43 Å of 330 SOSIP.664 Env trimer.

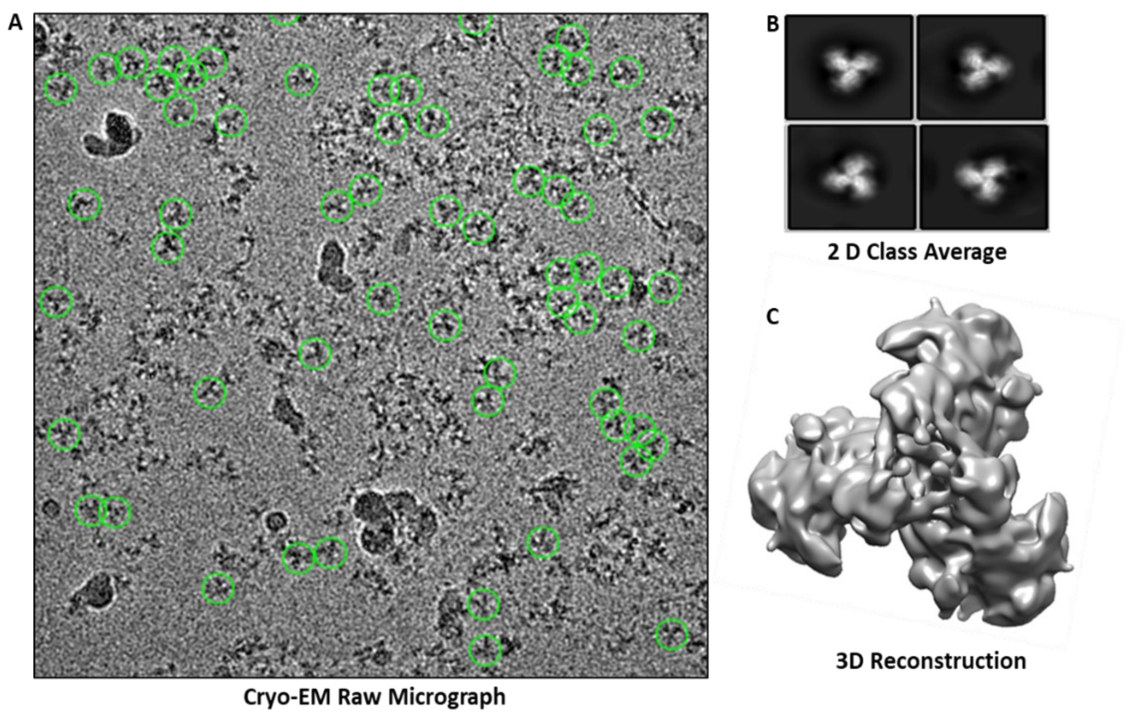

**Figure S7: Single particle cryo-Electron Microscopy (cryo-EM) of native like 330 SOSIP gp140 env trimers (A)** Raw Cryo-EM micrograph representing 330 SOSIP env trimers **(B)** Cryo-EM 2-D class averages of SOSIP gp140 env trimers **(C)** 3D reconstruction of soluble gp140 trimers in prefusion closed conformation.

Cryo-EM data processing pipeline of 330 Env complexed with 44m bnAb

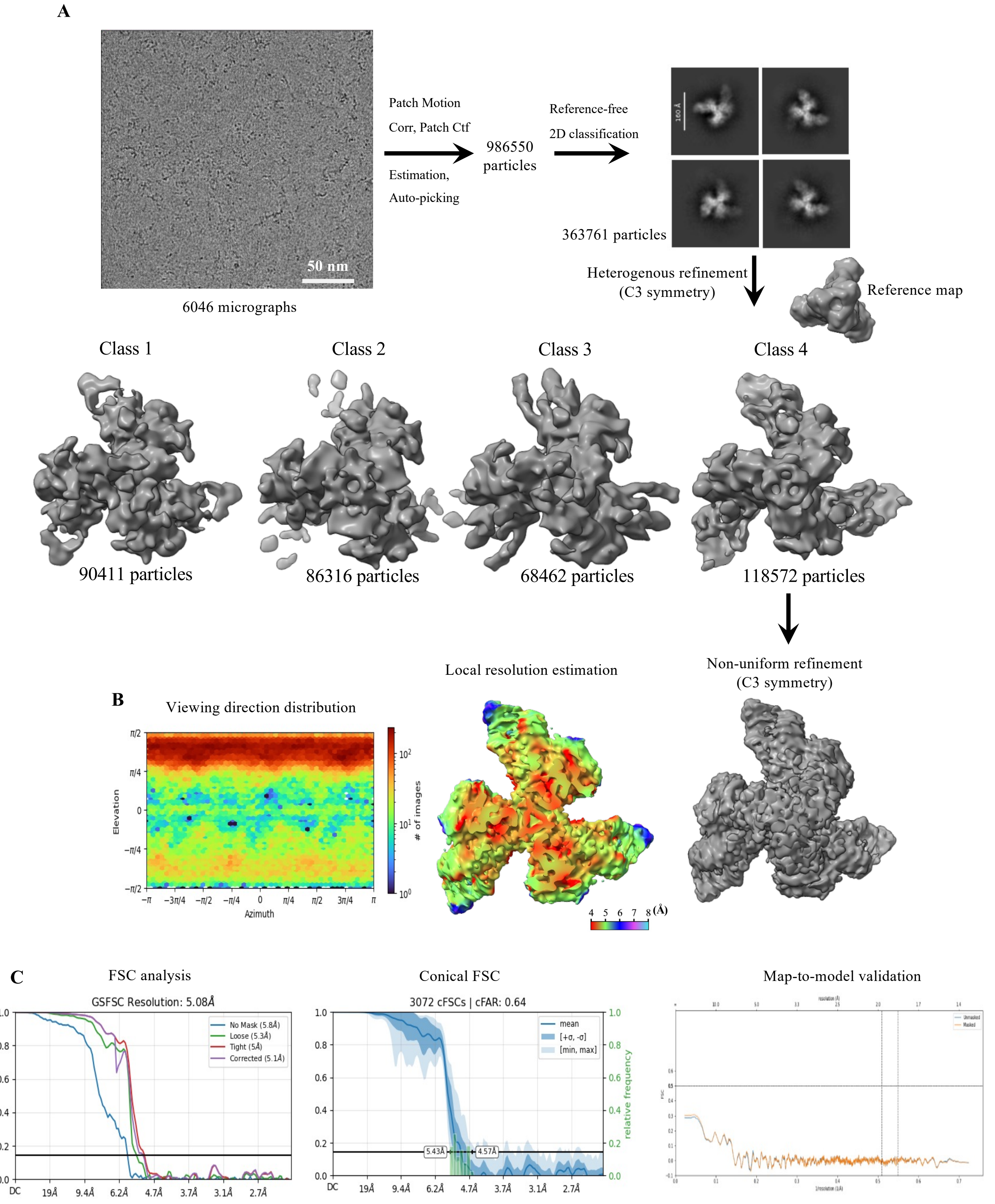

**Figure S8: Cryo-EM analysis of 330 SOSIP.664 Env trimer complexed with autologous bnAb 44m. (A)** Pipeline of Cryo-EM data processing using single particle cryo-EM and 3D classification of 330 SOSIP.664 Env trimer in complex with 44m bnAb. Ab initio reconstruction showed that the Class 04 map has better features with bnAb, and was therefore further considered for refinement and sharpening. **(B)** The viewing-direction particle distribution for Class 04 reveals diverse particle orientations in the dataset. **(C)** Gold standard Fourier Shell Correlation (FSC) curve of the cryo-EM map at 5.08 Å of 330 SOSIP Env trimer in complex with 44m bnAb.

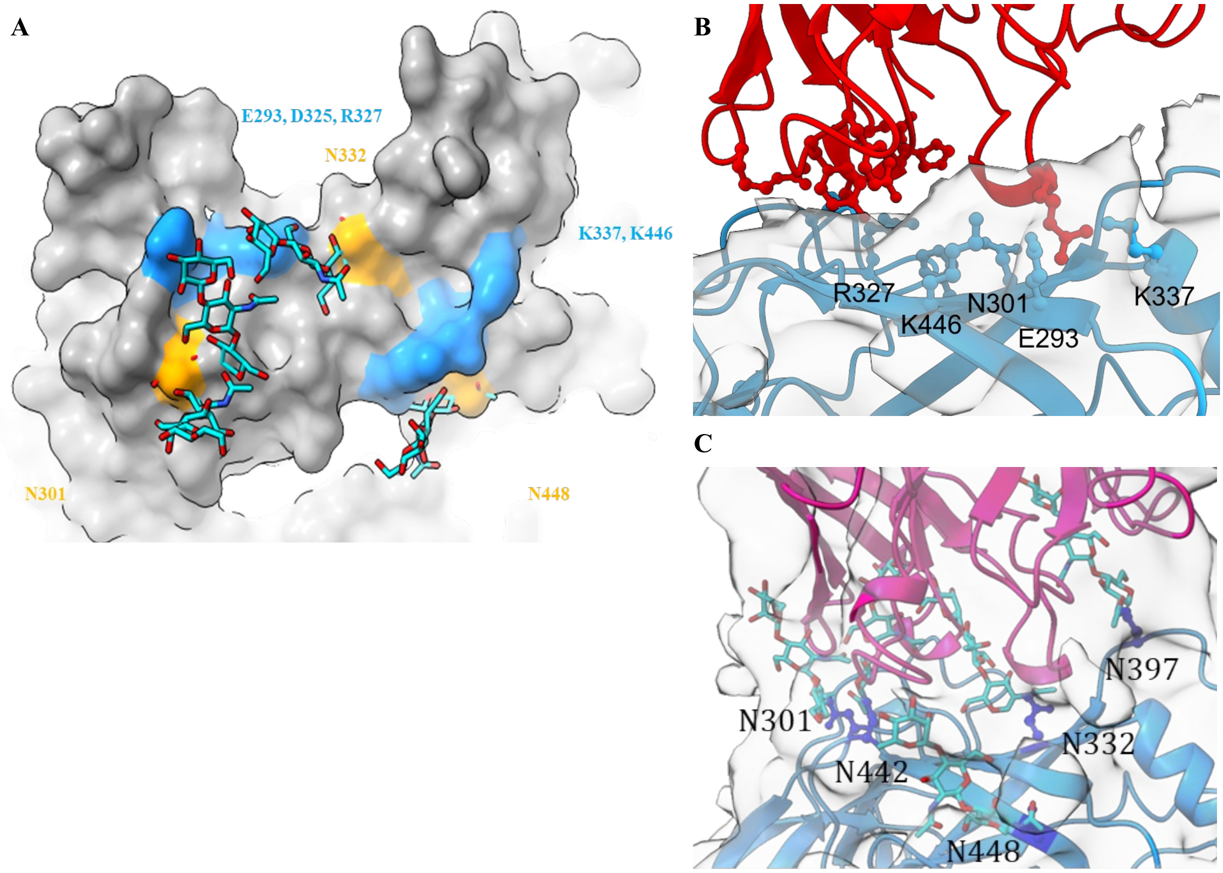

**Figure S9: Structural interface analysis of the autologous Env 330 SOSIP.664 in complex with V3-directed bnAb 44m (A)** Surface representation of gp120 residues and glycans involved in interactions with 44m. Contacting protein atoms are shown in blue, contacting glycan atoms in orange, and non-interacting regions in gray. **(B)** Representative EM density showing the fitting of interface amino acid residues and highlighting their contribution to the buried surface area analysis. **(C)** Fitting of Env glycans into the EM density map, supporting the buried surface area analysis

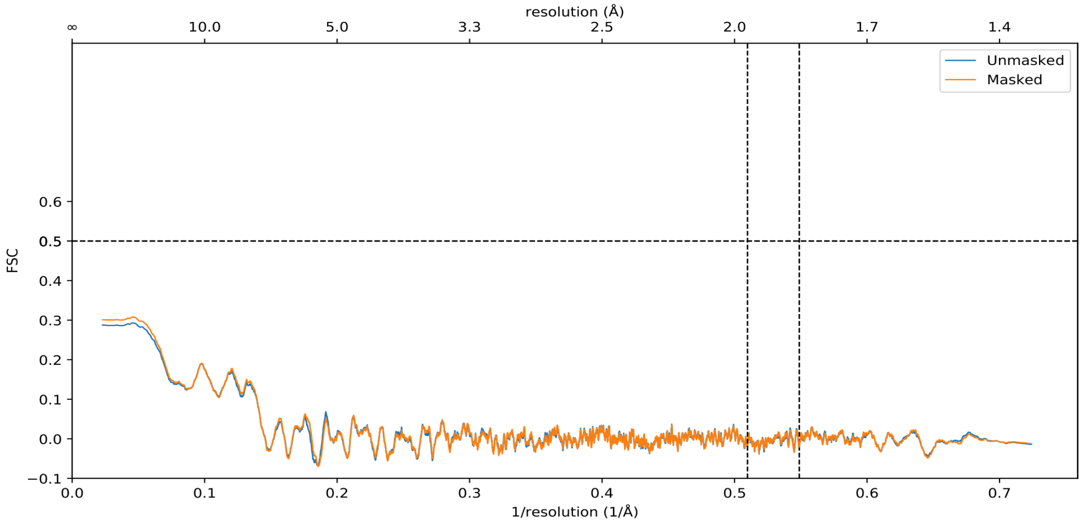

| | |
|---|---|
| **Validation** | |
| MolProbity Score | 2.52 |
| Clash Score | 17.36 |
| Rotamer Outliers (%) | 3.93 |
| Ramachandran Plot | |
| Outlier (%) | 0.09 |
| Allowed (%) | 4.59 |
| Favored (%) | 95.32 |

**Figure S10: Cryo-EM map-to-model validation of the AIIMS 330 Env-44m bnAb complex.** The AlphaFold-predicted models of AIIMS 330 Env and the autologous 44m bnAb were fitted into the 5.1 Å cryo-EM density map using ChimeraX and Coot. The fitted model was subsequently used for atomic model building and real-space refinement in Phenix 1.19.2. Map-to-model Fourier shell correlation (FSC) analysis was performed to assess the agreement between the refined atomic model and the cryo-EM density map. Refinement and model-validation statistics are summarized in **Table S8**.

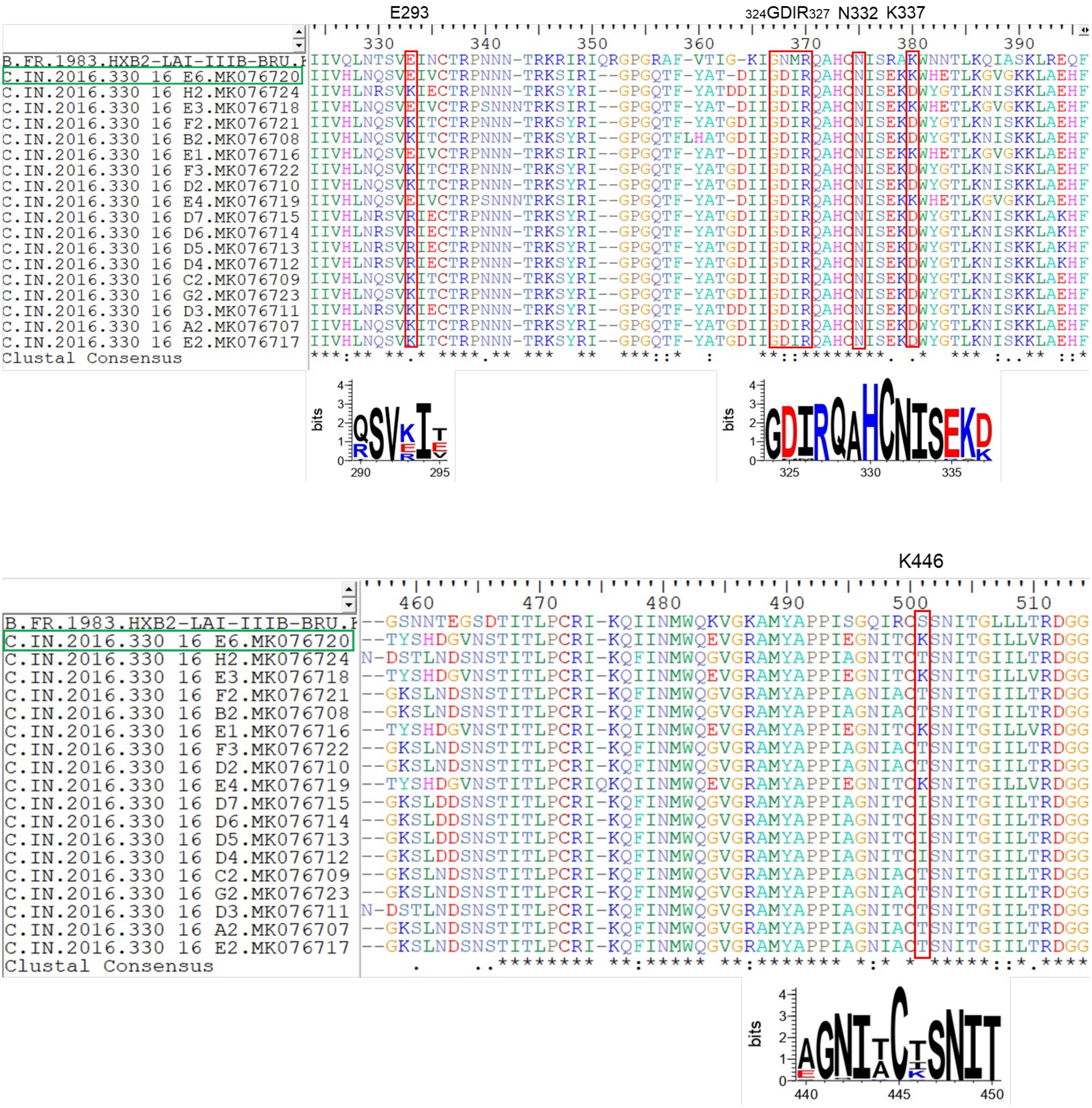

**Figure S11: Sequence alignment analysis of Env variants.** Sequence alignment of 18 HIV-1 Env variants generated from donor AIIMS 330 by single-genome amplification (SGA) from the previous study. The 330.E6.16 Env, used for immunogen design and structural characterization in complex with 44m by cryo-EM, is highlighted in green. Conserved V3-glycan supersite elements, including the N332 glycan and GDIR motif, are shared across all Env variants, whereas additional residues E293, K337, and K446 identified at the 330 Env-44m interface are restricted to E6, E1, E4, and E3 Env variants. Sequence conservation analysis and amino acid frequency at selected positions are highlighted using WebLogo.

**Table S1. Envelope trimer-stabilizing mutations were introduced into the 330 SOSIP and NFL constructs.**

| Stabilizing Mutations | 330_SOSIP | 330_NFL | Role in stabilization |
|---|---|---|---|
| [1]Δ MPER | + | + | Soluble prefusion conformation |
| [1]R6 Furin site | + | - | Soluble prefusion conformation |
| [1]I559P (IP) | + | + | Soluble prefusion conformation |
| [1]501C-605C (SOS) | + | - | Soluble prefusion conformation |
| [2](GGGGS)*2 linker | - | + | Soluble prefusion conformation |
| [3]65K | + | + | Reduced exposure for V3 non-nAbs |
| [3]535M | + | - | Reduced exposure for V3 non-nAbs |
| [3]543N | + | - | Reduced exposure for V3 non-nAbs |
| [4]201C and 433C | + | + | Prevent sCD4 exposure |
| [5]47D | + | + | BG505 Trimer Derived (TD8) stabilizing mutations |
| [5]49E | + | + | BG505 Trimer Derived (TD8) stabilizing mutations |
| [5]65K | + | + | BG505 Trimer Derived (TD8) stabilizing mutations |
| [5]106T | + | + | BG505 Trimer Derived (TD8) stabilizing mutations |
| [5]165L | + | + | BG505 Trimer Derived (TD8) stabilizing mutations |
| [5]429R | + | + | BG505 Trimer Derived (TD8) stabilizing mutations |
| [5]432Q | + | + | BG505 Trimer Derived (TD8) stabilizing mutations |
| [5]500R | + | + | BG505 Trimer Derived (TD8) stabilizing mutations |
| [6]171K | + | + | V2 Apex optimization |
| [7]519S/R | S | R | Increased expression and trimerization |
| [7]520L/R | L | R | Increased expression and trimerization |
| [7]568D/G | D | G | Increased expression and trimerization |
| [7]569G | - | + | Increased expression and trimerization |
| [7]570H | + | - | Increased expression and trimerization |
| [7]585H | + | - | Increased expression and trimerization |

[1] described in Sanders et. al. | 2013, PLoS Pathogens

[2] described in Sharma et. al. | 2015, Cell Reports

[3] described in Taeye et al., | 2015 | Cell

[4]described in Guenega et al. | 2017 | Immunity

[5]described in Guenega et al. | 2016 | JVI

[6]described in Andrabi et al. | 2015 | Immunity

[7]described in Steichen et al. | 2016 | Immunity

**Table S2. Binding kinetics of 330 Env Variants Assessed via Biolayer Interferometry.** A summarized table outlines the equilibrium dissociation constant (KD), association constant (Ka), and dissociation constant (Kdis) for bnAbs & non-nAbs tested with 330 gp140 envelope trimers (SOSIP & NFL) and ferritin NP's. The KD values were derived using 1:1 global fitting.

*NB- No Binding*

| mAbs | KD (nM) | KD Error | ka (1/Ms) | ka Error | kdis (1/s) | kdis Error | Full X^2 | Full R^2 |
|---|---|---|---|---|---|---|---|---|
| PGT145 | 1.070E-09 | 1.16E-11 | 9.520E05 | 6.325E03 | 1.019E-03 | 8.756E-06 | 30.4837 | 0.7208 |
| PGDM1400 | 1.344E-09 | 1.58E-11 | 4.031E05 | 1.638E03 | 5.417E-04 | 5.997E-06 | 17.0672 | 0.9131 |
| PGT151 | 5.009E-12 | - | 6.775E04 | 1.261E02 | 3.393E-07 | - | 7.6675 | 0.9914 |
| F105 | 7.158E-06 | 6.31E-05 | 4.048E02 | 3.568E03 | 2.897E-03 | 2.624E-04 | 17.3859 | - |
| 17b+sCD4 | NB | - | 7.747E04 | 4.022E03 | 2.441E-07 | - | 6.2921 | 0.3134 |
| | **SOSIP.664** | | | | | | | |

| mAbs | KD (nM) | KD Error | ka (1/Ms) | ka Error | kdis (1/s) | kdis Error | Full X^2 | Full R^2 |
|---|---|---|---|---|---|---|---|---|
| PGT145 | 1.39E-09 | 7.87E-12 | 4.344E05 | 1.854E03 | 6.045E-04 | 2.243E-06 | 15.3614 | 0.9917 |
| PGDM1400 | 3.32E-09 | 2.58E-11 | 1.783E05 | 9.826E02 | 5.926E-04 | 3.241E-06 | 7.1287 | 0.9874 |
| PGT151 | 1.42E-11 | - | 1.777E04 | 1.362E03 | 2.520E-07 | - | 0.9021 | 0.7821 |
| F105 | 2.35E-08 | 2.88E-10 | 7.727E04 | 8.742E02 | 1.812E-03 | 8.606E-06 | 0.5431 | 0.9632 |
| 17b+sCD4 | NB | - | 4.165E05 | 4.371E03 | 3.810E-07 | - | 0.1097 | 0.9624 |
| | **NFL.664** | | | | | | | |

| mAbs | KD (nM) | KD Error | ka (1/Ms) | ka Error | kdis (1/s) | kdis Error | Full X^2 | Full R^2 |
|---|---|---|---|---|---|---|---|---|
| PGT145 | 1.1E-09 | 7.93E-12 | 7.139E05 | 2.550E03 | 7.840E-04 | 4.920E-06 | 46.0383 | 0.9144 |
| PGDM1400 | 1.39E-09 | 1.99E-11 | 3.030E05 | 1.164E03 | 4.211E-04 | 5.811E-06 | 24.6365 | 0.9285 |
| PGT151 | 3.02E-12 | - | 6.651E04 | 1.488E02 | 2.010E-07 | - | 8.2643 | 0.9862 |
| F105 | 6.99E-08 | 1.76E-09 | 3.144E04 | 6.405E02 | 2.198E-03 | 3.229E-05 | 2.1896 | 0.7236 |
| 17b+sCD4 | NB | - | 1.233E05 | 3.768E03 | 2.441E-07 | - | 1.3081 | 0.3051 |
| | **Ferritin NP** | | | | | | | |

**Table S3: Comparison of potential N-linked glycosylation sites (PNGS) across cross-clade envelopes.** An N indicates the presence of a PNGS, if NXT/S motif in a.a. sequence, whereas X should not be proline. BG505 & 330 Envs share the same total number of PNGS and the same number of NXT & NXS N-glycosylation sites. Created by the N-glycosite tool using the Los Alamos HIV Sequence Database

| Envelope | PNGS | NXT | NXS | 88 | 133 | 135 | 136 | 137 | 139 | 141 | 142 | 147 | 156 | 160 | 186 | 187 | 188 | 197 | 230 | 234 | 241 | 262 | 276 | 289 | 295 | 301 | 332 |
|---|---|---|---|---|---|---|---|---|---|---|---|---|---|---|---|---|---|---|---|---|---|---|---|---|---|---|---|
| BG505 | 28 | 16 | 12 | N | N | - | N | - | - | N | N | - | N | N | N | - | - | N | N | N | N | N | N | N | N | N | N |
| 330 | 28 | 16 | 12 | N | - | - | N | - | - | N | - | - | N | N | N | N | - | N | N | N | N | N | N | N | N | N | N |
| 16055 | 28 | 17 | 11 | N | - | - | N | - | N | N | - | - | N | N | N | - | - | N | N | N | N | N | N | N | N | N | N |
| C97ZA | 30 | 20 | 10 | N | N | - | N | N | - | N | - | - | N | N | N | N | - | N | N | N | N | N | N | N | N | N | N |
| 42J41 | 29 | 21 | 8 | N | - | - | N | - | - | N | - | - | N | N | N | N | - | N | N | N | N | N | N | N | N | N | - |
| Du422 | 30 | 19 | 11 | N | - | N | N | - | - | N | - | N | N | N | N | - | - | N | N | N | N | N | N | N | N | N | N |
| ZM246F | 29 | 18 | 11 | N | - | - | N | - | - | N | - | - | N | N | N | N | N | N | N | N | N | N | N | N | N | N | N |

| Envelope | PNGS | NXT | NXS | 334 | 339 | 355 | 356 | 360 | 363 | 386 | 392 | 397 | 398 | 399 | 403 | 404 | 406 | 407 | 409 | 411 | 442 | 448 | 461 | 462 | 463 | 505 | 611 | 616 | 618 | 624 | 625 | 637 |
|---|---|---|---|---|---|---|---|---|---|---|---|---|---|---|---|---|---|---|---|---|---|---|---|---|---|---|---|---|---|---|---|---|
| BG505 | 28 | 16 | 12 | - | N | N | N | - | N | N | N | N | N | - | - | - | N | - | - | N | - | N | - | N | N | - | N | N | N | N | N | N |
| 330 | 28 | 16 | 12 | - | N | N | N | - | - | N | N | N | - | - | - | - | N | N | - | N | N | N | - | N | N | - | N | N | - | N | N | N |
| 16055 | 28 | 17 | 11 | - | N | - | N | N | - | N | N | N | - | N | - | N | N | - | N | - | N | N | - | - | N | - | N | N | - | N | N | N |
| C97ZA | 30 | 20 | 10 | - | N | - | N | - | N | N | N | N | - | - | - | - | N | - | - | N | N | N | - | - | N | N | N | N | - | N | N | N |
| 42J41 | 29 | 21 | 8 | N | N | N | N | - | - | N | N | N | - | - | N | N | N | - | - | N | N | N | - | N | N | - | N | N | N | N | - | N |
| Du422 | 30 | 19 | 11 | - | N | - | N | - | - | N | N | N | - | - | - | - | N | N | - | - | N | N | N | - | N | - | N | N | - | N | N | N |
| ZM246F | 29 | 18 | 11 | - | N | - | N | - | - | N | N | N | - | - | - | - | N | N | - | N | N | N | N | - | N | - | N | N | - | N | N | N |

**Table S4. Mutant pseudovirus neutralization assay to assess residue dependence.** Cryo-EM identified Env contact residues (E293, D325, R327, H330, N332, K337, and K446) were individually mutated by site-directed mutagenesis (alanine substitutions, except N332Q). Mutant pseudoviruses were tested for antibody neutralization using a TZM-bl reporter assay. $IC_{50}$ values represent the mean of independent experiments.

| Mutant Neutralization | | |
|---|---|---|
| IC50 µg/mL | | Fold Change |
| RTE6 WT | 0.086 | 1 |
| E293A | 0.065 | 0.76 |
| D325A | 0.041 | 0.48 |
| R327A | >10 | 116 |
| H330A | 0.092 | 1.07 |
| N332Q | >10 | 116 |
| K337A | 0.055 | 0.64 |
| K446A | <0.04 | 0.47 |

**Table S5: Midpoint neutralization titers of rabbit immune sera at week 22 tested against a panel of Env-pseudotyped viruses.** TZM-bl neutralization assays were performed in duplicate, and MLV (murine leukemia virus) was used as a negative control. The study is divided into four groups, Group 1, 2 & 3 (n=4) represent 330 NFL, SOSIP trimers & multivalent Ferritin NP's, while Group 4 (n=2) represents a placebo. The $ID_{50}$ values, i.e., the serum dilution at which infectivity was inhibited by 50%, are shown and color-coded: white = no neutralization, $ID_{50}$ < 20; yellow = weak neutralization, $ID_{50}$ 20-100; orange = moderate neutralization, 100-300; red = strong neutralization, 300-2000. Individual $ID_{50}$ value of immunized animal tested against env pseudotyped viruses.

| | | | **virus** | **MLV** | **330.16.E6** | **329.14.B1** | **MW965.26** | **25925** | **SF162** | **Bal01** | **BG505** |
|---|---|---|---|---|---|---|---|---|---|---|---|
| | | | **Tier** | **N/A** | **ND** | **ND** | **1A** | **1B** | **1A** | **1A** | **2** |
| **Study** | **Groups** | **Immunogen** | **Clade** | **control** | **C** | **C** | **C** | **C** | **B** | **B** | **A** |
| **LBPL/NG-1736 (EF)** | Group 1 | **Ferritin NP** | **NZW5265** | <20 | 1230 | 410 | 277 | 100 | 100 | <20 | <20 |
| | | | **NZW5266** | <20 | 1450 | 530 | 255 | 77 | 122 | <20 | <20 |
| | | | **NZW5267** | <20 | 1090 | 450 | 344 | 77 | 122 | <20 | <20 |
| | | | **NZW5268** | <20 | 677 | 33 | 211 | 33 | 63 | <20 | <20 |
| | Group 2 | **SOSIP** | **NZW5269** | <20 | 890 | 470 | 100 | 33 | 55 | <20 | <20 |
| | | | **NZW5270** | <20 | 830 | 473 | 166 | <20 | 100 | 32 | <20 |
| | | | **NZW5271** | <20 | 710 | 450 | 144 | 100 | 144 | <20 | <20 |
| | | | **NZW5272** | <20 | 830 | 270 | 144 | 77 | 77 | <20 | <20 |
| | Group 3 | **NFL** | **NZW5273** | <20 | <20 | <20 | <20 | <20 | <20 | <20 | <20 |
| | | | **NZW5274** | <20 | 155 | 60 | 36 | 90 | <20 | <20 | <20 |
| | | | **NZW5275** | <20 | 375 | 232 | 73 | 96 | 25 | <20 | <20 |
| | | | **NZW5276** | <20 | 429 | 393 | 41 | 65 | <20 | <20 | <20 |
| | Group 4 | **Placebo** | **NZW5277** | <20 | <20 | <20 | <20 | <20 | <20 | <20 | <20 |

| **ID50** |
|---|
| <20 |
| 20-100 |
| 100-300 |
| 300-2000 |

**Table S6: BLI-Octet tandem competition matrix.** Heatmap illustrating competitive binding between immune sera and reference HIV-1 bnAbs and non-neutralizing antibodies. A 35% competition threshold was used to define competing interactions.

Competition Matrix

| Immune Sera | Ab# | PGT145 | PGDM1400 | VRC01 | 44m | 10.1074 | PGT121 | F105 | 447-52d | 412d | 17b | PGT151 |
|---|---|---|---|---|---|---|---|---|---|---|---|---|
| SOSIP | Placebo W0 | 0.1231 | 0.1192 | 0.0337 | 0.1703 | 0.1535 | 0.1428 | 0.1042 | 0.1623 | 0.0755 | 0.0894 | 0.3663 |
| SOSIP | Placebo W22 | 0.1992 | 0.2432 | 0.0897 | 0.3465 | 0.2895 | 0.2018 | 0.1569 | 0.1832 | 0.1295 | 0.1102 | 0.2986 |
| SOSIP | Ferritin NP W0 | 0.2388 | 0.2814 | 0.1244 | 0.3688 | 0.3207 | 0.2330 | 0.2160 | 0.2632 | 0.2000 | 0.1632 | 0.3601 |
| SOSIP | Ferritin NP W22 | 0.0306 | 0.0465 | 0.0109 | -0.0155 | 0.0342 | 0.0768 | 0.0845 | 0.1714 | 0.1281 | 0.0783 | 0.2296 |
| SOSIP | SOSIP W0 | 0.1697 | 0.1869 | 0.0683 | 0.1748 | 0.1919 | 0.1643 | 0.1664 | 0.2584 | 0.1908 | 0.1199 | 0.3947 |
| SOSIP | SOSIP W22 | -0.0597 | -0.0358 | -0.0792 | -0.0955 | -0.0344 | -0.0352 | 0.0443 | 0.1607 | 0.0977 | 0.0384 | 0.2173 |
| NFL | Placebo W0 | | 0.1114 | 0.0013 | 0.1198 | 0.1615 | 0.1693 | 0.1666 | 0.2377 | 0.2341 | 0.1860 | 0.2986 |
| NFL | Placebo W22 | 0.1609 | 0.1794 | 0.0770 | 0.2573 | 0.2952 | 0.1830 | 0.1891 | 0.2603 | 0.2452 | 0.1684 | 0.3663 |
| NFL | NFLW0 | 0.1966 | 0.2352 | 0.1292 | 0.3063 | 0.3546 | 0.2270 | 0.0813 | 0.1318 | 0.1224 | 0.0735 | 0.2093 |
| NFL | NFLW22 | -0.1333 | -0.1297 | -0.1213 | -0.1239 | -0.0343 | 0.0045 | -0.0157 | 0.0660 | 0.0250 | -0.0396 | 0.2296 |

**Table S7: Cryo-EM data collection and processing of unliganded 330 SOSIP.664 Env trimer.**

| Data collection and processing | |
|---|---|
| Magnification | 42000 |
| Voltage (kV) | 200 |
| Electron exposure (e–/Å2) | 60 |
| Number of frames | 20 |
| Defocus range (μm) | -0.75 to –2.25 |
| Pixel size (Å) | 1.17 |
| Symmetry imposed | C3 |
| Number of movies | 4353 |
| Number of particles in model | 89209 |
| Map resolution (Å) | 7.43 |
| FSC threshold | 0.143 |

**Table S8: Cryo-EM data collection and processing of Env 330-44m bnAb complex.**

| **Data collection and processing** | |
|---|---|
| Magnification | 42000 |
| Voltage (kV) | 200 |
| Electron exposure (e–/Å2) | 60 |
| Number of frames | 20 |
| Defocus range (μm) | -0.75 to –2.25 |
| Pixel size (Å) | 1.17 |
| Symmetry imposed | C3 |
| Number of movies | 6046 |
| Number of particles in model | 118572 |
| Map resolution (Å) | 5.1 |
| FSC threshold | 0.143 |
| **Validation** | |
| MolProbity Score | 2.52 |
| Clash Score | 17.36 |
| Rotamer Outliers (%) | 3.93 |
| Ramachandran Plot | |
| Outlier (%) | 0.09 |
| Allowed (%) | 4.59 |
| Favored (%) | 95.32 |

**Table S9: Comparison of rabbit serum autologous neutralization titers elicited by different Env SOSIP trimers.** Neutralization titers are presented as reciprocal serum dilutions corresponding to 50% inhibition $ID_{50}$ of pseudoviruses expressing autologous Envs in TZM-bl cells.

| | SOSIP.664 Env trimers | | | | | | | | |
|---|---|---|---|---|---|---|---|---|---|
| Envelope | **330** | **329** | **ZM197** | **IPGE-THIVC** | **AMC009** | **B41** | **AMC008** | **BG505 (v4.2)** | **BG505 (v5.2)** |
| Subtypes | C | C | C | C | B | B | B | A | A |
| Origin | Pediatric | Pediatric | Adult | Adult | | Adult | Adult | Pediatric | Pediatric |
| Categorization | Elite neutralizer | Elite neutralizer | Elite neutralizer | Elite neutralizer | Elite neutralizer | Elite neutralizer | Elite neutralizer | Founder virus | Founder virus |
| Disease Stage | Late Stage | Late Stage | Late Stage | Late Stage | Early Stage | Late Stage | Early Stage | Early Stage | Early Stage |
| Animal No. | Neutralization Titers ($ID_{50}$) | | | | | | | | |
| 1 | 890 | 38 | 1010 | 2284 | <20 | 249 | 219 | 1190 | >4860 |
| 2 | 830 | 58 | 153 | 431 | <20 | 965 | 299 | 2022 | 2554 |
| 3 | 710 | 50 | 64 | 104 | <20 | 111 | 262 | 627 | 50 |
| 4 | 830 | 58 | 45 | 342 | <20 | 601 | 272 | 1556 | <20 |
| 5 | – | – | 114 | – | 48 | <20 | 178 | 738 | 494 |
| Average | 815 | 51 | 277 | 790 | 48 | 481.5 | 246 | 1226.6 | 1990 |

| | | | | | | | | | |
|---|---|---|---|---|---|---|---|---|---|
| References | Current study | S. Kumar et al \| vaccines | Torrents et al. \| JVI | R. Kumar et al. \| PLOS pathogens | Schorcht et al. \| JVI | de Taeye et al.\| Cell | Torrents et al. \| JVI | de Taeye et al.\| Cell | Torrents et al. \| JVI |

**Table S10. Longitudinal Binding Kinetics, Durability, and Boosting Profiles across Autologous and Heterologous Viral Strains.** Data values represent mean binding levels ± standard deviation (SD). *P < 0.05, **P < 0.01 indicate statistically significant differences in cross-sectional post-hoc Tukey HSD pairwise multi-comparisons among the active immunogen variants. Global Linear Mixed-Effects Models (LMM) demonstrated highly significant main group effects, time effects, and group × time interactions across all tested viral populations (P < 0.001).

| Virus / Strain | Immunogen Group | Baseline (W0) | Peak Boost 1 (W6) | Durability Trough (W20) | Recall Boost 2 (W22) | Statistical Comparisons (Tukey HSD Post-Hoc) |
|---|---|---|---|---|---|---|
| **330 (Autologous)** | 330 Ferritin NP (n=4) | 0.273 ± 0.026 | 1.539 ± 0.151 | 0.488 ± 0.100 | 1.874 ± 0.054 | Global LMM: Group, Time, Group×Time: all P < 0.001. W6: NP vs SOSIP (P=0.681), NP vs NFL (P=0.019)*, SOSIP vs NFL (P=0.048)* W20: NP vs SOSIP (P=0.035)*, NP vs NFL (P=0.041)*, SOSIP vs NFL (P=0.002)** W22: NP vs SOSIP (P=0.002)**, NP vs NFL (P=0.001)**, SOSIP vs NFL (P=0.314) |
| | 330 SOSIP (n=4) | 0.268 ± 0.096 | 1.453 ± 0.147 | 0.664 ± 0.158 | 1.497 ± 0.227 | |
| | 330 NFL (n=4) | 0.315 ± 0.073 | 1.209 ± 0.059 | 0.251 ± 0.106 | 1.317 ± 0.076 | |
| | Placebo (n=2) | 0.030 ± 0.002 | 0.070 ± 0.003 | 0.084 ± 0.007 | 0.066 ± 0.005 | |
| **SF162 (Heterologous)** | 330 Ferritin NP (n=4) | 0.091 ± 0.067 | 1.428 ± 0.104 | 0.096 ± 0.029 | 1.669 ± 0.042 | Global LMM: Group, Time, Group×Time: all P < 0.001. W6: NP vs SOSIP (P=0.851), NP vs NFL (P=0.900), SOSIP vs NFL (P=0.627) W20: NP vs SOSIP (P=0.278), NP vs NFL (P=0.900), SOSIP vs NFL (P=0.188) W22: NP vs SOSIP (P=0.677), NP vs NFL (P=0.900), SOSIP vs NFL (P=0.746) |
| | 330 SOSIP (n=4) | 0.062 ± 0.062 | 1.477 ± 0.119 | 0.422 ± 0.485 | 1.496 ± 0.499 | |
| | 330 NFL (n=4) | 0.039 ± 0.020 | 1.390 ± 0.164 | 0.040 ± 0.028 | 1.644 ± 0.035 | |
| | Placebo (n=2) | 0.030 ± 0.002 | 0.070 ± 0.003 | 0.084 ± 0.007 | 0.066 ± 0.005 | |
| **6240 (Heterologous)** | 330 Ferritin NP (n=4) | 0.079 ± 0.052 | 1.524 ± 0.085 | 0.068 ± 0.025 | 1.640 ± 0.071 | Global LMM: Group, Time, Group×Time: all P < 0.001. W6: NP vs SOSIP (P=0.015)*, NP vs NFL (P=0.210), SOSIP vs NFL (P=0.248) W20: NP vs SOSIP (P=0.239), NP vs NFL (P=0.655), SOSIP vs NFL (P=0.063) W22: NP vs SOSIP (P=0.112), NP vs NFL (P=0.294), SOSIP vs NFL (P=0.774) |
| | 330 SOSIP (n=4) | 0.043 ± 0.030 | 1.334 ± 0.050 | 0.106 ± 0.041 | 1.534 ± 0.048 | |
| | 330 NFL (n=4) | 0.037 ± 0.018 | 1.425 ± 0.085 | 0.048 ± 0.023 | 1.565 ± 0.074 | |
| | Placebo (n=2) | 0.030 ± 0.002 | 0.070 ± 0.003 | 0.079 ± 0.001 | 0.056 ± 0.005 | |
| **1086 (Heterologous)** | 330 Ferritin NP (n=4) | 0.072 ± 0.052 | 1.067 ± 0.104 | 0.079 ± 0.020 | 1.271 ± 0.195 | Global LMM: Group, Time, Group×Time: all P < 0.001. W6: NP vs SOSIP (P=0.212), NP vs NFL (P=0.002)**, SOSIP vs NFL (P=0.026)* W20: NP vs SOSIP (P=0.494), NP vs NFL (P=0.758), SOSIP vs NFL (P=0.200) W22: NP vs SOSIP (P=0.844), NP vs NFL (P=0.050)*, SOSIP vs NFL (P=0.117) |
| | 330 SOSIP (n=4) | 0.068 ± 0.029 | 0.091 ± 0.174 | 0.148 ± 0.143 | 1.225 ± 0.054 | |
| | 330 NFL (n=4) | 0.029 ± 0.013 | 0.646 ± 0.027 | 0.038 ± 0.009 | 1.037 ± 0.037 | |
| | Placebo (n=2) | 0.030 ± 0.002 | 0.070 ± 0.003 | 0.084 ± 0.007 | 0.066 ± 0.005 | |
| **329 (Heterologous)** | 330 Ferritin NP (n=4) | 0.051 ± 0.016 | 1.456 ± 0.055 | 0.062 ± 0.026 | 1.458 ± 0.115 | Global LMM: Group, Time, Group×Time: all P < 0.001. W6: NP vs SOSIP (P=0.658), NP vs NFL (P=0.784), SOSIP vs NFL (P=0.900) W20: NP vs SOSIP (P=0.383), NP vs NFL (P=0.900), SOSIP vs NFL (P=0.420) W22: NP vs SOSIP (P=0.031)*, NP vs NFL (P=0.087), SOSIP vs NFL (P=0.775) |
| | 330 SOSIP (n=4) | 0.099 ± 0.064 | 1.513 ± 0.138 | 0.223 ± 0.281 | 1.672 ± 0.107 | |
| | 330 NFL (n=4) | 0.064 ± 0.015 | 1.498 ± 0.057 | 0.071 ± 0.014 | 1.626 ± 0.062 | |
| | Placebo (n=2) | 0.059 ± 0.015 | 0.082 ± 0.010 | 0.149 ± 0.012 | 0.075 ± 0.004 | |
